# Depletion of LKB1 in cerebellar granule cells affects parallel fiber–Purkinje cell synaptic organization and causes progressive locomotion deficits

**DOI:** 10.64898/2026.11.16.766918

**Authors:** Ryo Kozawa, Sota Sawada, Hinako Kiumi, Shun Hamada, Mayu Kitamura, Masataka Yamamoto, Mayumi Wakazaki, Mayuko Sato, Kiminori Toyooka, Nobuhiko Ohno, Hiroshi Takemura, Toshihisa Ohtsuka, Akari Hagiwara

## Abstract

The cerebellum is the center of motor coordination and learning, where it integrates sensory and motor information and transmits signals to the muscles via motor neurons. The sole cerebellar output neurons, Purkinje cells (PCs), receive two main inputs from climbing fibers (CFs) and parallel fibers (PFs), which are the axons of granule cells (GCs). The serine/threonine kinase liver kinase B1 (LKB1) regulates GC migration; however, its role in PF synapses remains unclear. We generated a novel strain of mice with a LKB1 conditional knockout (cKO) in GCs during the postnatal period to investigate the presynaptic function of LKB1. LKB1 cKO mice exhibited progressive declines in motor coordination, whereas general locomotor activity was preserved. Morphological analysis revealed shrinkage of the cerebellar molecular layer with increased microglial density, suggesting degenerative remodeling of PF–PC synapses. Conventional electron microscopy showed a slight increase in the PF terminal area in LKB1 cKO mice, whereas three-dimensional reconstructions revealed no significant differences in bouton morphology except for an increased mitochondrial volume ratio. In contrast, the proportion of PF boutons forming multiple synapses with PC spines was higher in LKB1 cKO mice than in controls, potentially altering synaptic inputs to PCs and contributing to downstream changes in cerebellar output and reduced presynaptic occupancy at neuromuscular junctions. These findings highlight the importance of LKB1 in GCs for maintaining PF–PC synaptic remodeling and cerebellar circuit integrity, which may contribute to motor coordination.

## INTRODUCTION

The development and maintenance of neural circuits involve diverse intracellular signaling pathways that regulate neuronal morphogenesis and synaptic organization. The ubiquitously expressed serine/threonine kinase liver kinase B1 (LKB1, also known as STK11) serves as a master regulator of cellular proliferation, energy homeostasis, and neuronal morphogenesis. LKB1 activates multiple downstream kinases of the AMPK subfamily, including AMPKα1/2, NUAKs, SAD-A/B, and MARKs, to regulate diverse cellular processes (Barnes et al. 2007; Courchet et al. 2013; Kuwako and Okano 2018a, 2018b). Among these pathways, LKB1–SAD-A/B signaling is required for axon specification and neuronal polarization in cortical neurons (Barnes et al. 2007; Shelly et al. 2007; Shelly and Poo 2011). SAD-B also regulates presynaptic vesicle dynamics and short-term synaptic plasticity through CAST phosphorylation and contributes to hippocampus-dependent memory (Mochida et al. 2016; Watabe et al. 2016). Furthermore, LKB1–AMPK signaling promotes PGC-1α-dependent mitochondrial biogenesis to support axonal growth, while the LKB1–NUAK1 pathway regulates mitochondrial capture at presynaptic sites and promotes terminal axon branching (Courchet et al. 2013; Vaarmann et al. 2016).

The cerebellum coordinates movements by integrating motor commands from the cerebral cortex with sensory feedback to ensure smooth and accurate movements. The primary output neurons of the cerebellum, Purkinje cells (PCs), receive excitatory inputs from granule cell (GC) axons, known as parallel fibers (PFs), and climbing fibers (CFs) from the inferior olive. These inputs converge onto PC dendritic spines within the molecular layer (ML). Cerebellar development begins during embryogenesis and involves the coordinated proliferation, migration, and differentiation of GCs and PCs. During embryonic development, PCs are generated between E10 and E13 and migrate into the developing cerebellum, while granule cell progenitors (GCPs) migrate from the rhombic lip to form the external granule layer (EGL) around E15, where they proliferate extensively during early postnatal development (Rahimi-Balaei et al. 2018). As GCs differentiate, they extend axons bidirectionally to form PFs, and their cell bodies migrate inward along Bergmann glial fibers to establish the granule cell layer (GL). Concurrently, PC dendrites extend into the ML, where they form synaptic connections with PFs. Previous studies have demonstrated that LKB1 plays an important role in the development of both GCs and PCs. In PCs, LKB1 regulates dendritic patterning, particularly dendritic self-avoidance, through SIK1/2-dependent regulation of Robo2 localization, which prevents inappropriate overlap between dendritic branches (Gibson et al. 2014; Kuwako and Okano 2018a, 2018b). However, embryonic deletion of LKB1 in GCs using Atoh1-Cre led to disrupted GC migration, excessive cerebellar foliation with enlargement, and severe motor coordination deficits in the rotarod test (Men et al. 2015; Ryan et al. 2017). Together, these findings established LKB1 as an important regulator of cerebellar development and neuronal morphogenesis. However, its role in the structural organization and maintenance of presynaptic terminals at PF–PC synapses remains unclear. In particular, whether LKB1 contributes to presynaptic organization beyond its established role in early GC development has not been fully investigated.

In this study, we crossed LKB1-floxed mice with E3CreN mice, in which Cre expression is driven by a downstream regulatory region of *Grin2C* (encoding the NMDA receptor subunit) to generate mice with postnatal LKB1 deletion in cerebellar GCs during synaptogenesis (Miyazaki et al. 2012); these mice are referred to herein as LKB1 conditional knockout (cKO) mice. Unlike embryonic LKB1 deletion in GCPs, which leads to abnormal cerebellar morphology and early-onset motor deficits by 4–5 weeks (Men et al. 2015; Ryan et al. 2017), postnatal LKB1 deletion showed less marked changes. However, beginning at around 10 weeks, LKB1 cKO mice developed progressive motor coordination impairments correlating with age-dependent cerebellar ML degeneration. This was accompanied by multisynaptic PF–PC connections. Furthermore, LKB1 cKO mice exhibited significant reductions in neuromuscular junction (NMJ) innervation, indicating compromised motor neuron transmission. These findings highlight a critical role for LKB1 in presynaptic remodeling and maintenance, with broad implications for cerebellar integrity and motor function.

## RESULTS

### Generation of LKB1 cKO in cerebellar granule cells during postnatal development

To investigate the role of LKB1 at PF–PC synapses, we generated LKB1 cKO mice using Grin2C-iCre to deplete LKB1 in cerebellar GCs during the postnatal period of synaptogenesis (Fig. 1A) (Miyazaki et al. 2012; Nakada, Saunders, and Morrison 2010). When Grin2C-iCre mice were crossed with Ai14 reporter mice, designed for tdTomato expression upon Cre-mediated recombination, tdTomato fluorescence was first detected in the posterior cerebellum at P3 and had expanded throughout the cerebellum by P21 (Fig. S1A).

**Figure 1.**
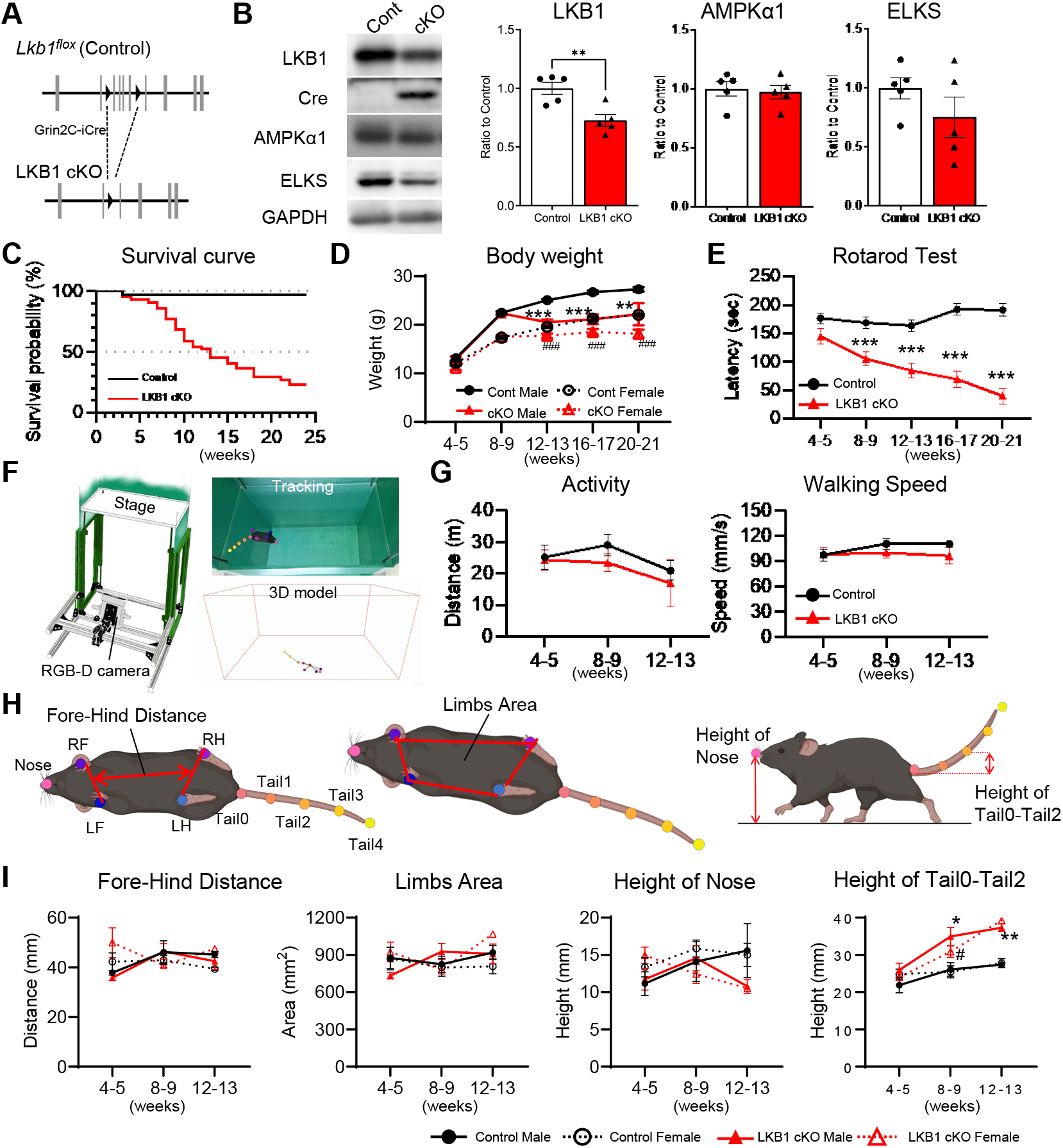
Characterization of the LKB1 cKO in cerebellar granule cells. (A) Schematic of the targeting strategy for cerebellar granule cell-specific deletion of *Lkb1*. Exons 3–6 of the LKB1 were deleted. (B) Immunoblot analysis of cerebellar lysates using indicated antibodies. LKB1 expression was significantly reduced following Cre recombinase expression in the LKB1 cKO mice at 5 weeks. Signal intensities were normalized to the control signal of each preparation. Control n = 5, LKB1 cKO n = 6, Mean ± SEM, ** p < 0.01, Student’s t-test. (C) Survival analysis showing that approximately 50% of LKB1 cKO mice died by 10 weeks. Control n = 161 (censored = 153, death = 8), LKB1 cKO n = 139 (censored = 74, death = 65). Gehan–Breslow– Wilcoxon test, p < 0.0001. Kaplan–Meier curve ± 95% CI. (D) Body weights across development were recorded in control and LKB1 cKO mice. Control male; 4–5 weeks = 25, 8–9 weeks = 36, 12–13 weeks = 35, 16–17 weeks = 37, 20–21 weeks = 26, Control female; 4–5 weeks = 23, 8–9 weeks = 27, 12–13 weeks = 30, 16–17 weeks = 27, 20–21 weeks = 27, LKB1 cKO male; 4–5 weeks = 23, 8–9 weeks = 19, 12–13 weeks = 23, 16–17 weeks = 9, 20–21 weeks = 6, LKB1 cKO female; 4–5 weeks = 21, 8–9 weeks = 29, 12–13 weeks = 21, 16–17 weeks = 12, 20–21 weeks = 11. (E) Motor coordination assessed using the accelerating rotarod. While control mice maintained performance with age, LKB1 cKO mice showed significantly reduced latency to fall in older age groups. Control; 4–5 weeks = 48, 8–9 weeks = 63, 12–13 weeks = 65, 16–17 weeks = 54, LKB1 cKO; 4–5 weeks = 44, 8–9 weeks = 48, 12–13 weeks = 44, 16–17 weeks = 21, 20–21 weeks = 17. Mean ± SEM. (F-G) Gait analysis using reconstructed 3D models to evaluate locomotor activity and gait parameters during walking. (H-I) No significant differences were observed in activity level, while LKB1 cKO mice displayed slightly elevated tail posture. Control; 4–5 weeks = 7 (male 2, female 5), 8–9 weeks = 8 (male 3, female 5), 12–13 weeks = 8 (male 3, female 5), LKB1 cKO; 4–5 weeks = 11 (male 7, female 4), 8–9 weeks = 9 (male 5, female 4), 12–13 weeks = 3 (male 2, female 1). Mean ± SEM, * p < 0.05, ** p < 0.01, *** p < 0.001. When data were analyzed separately by sex, significant differences in females are indicated by # p < 0.05, ## p < 0.01, ### p < 0.001, one-way ANOVA followed by Tukey’s multiple comparisons test.

In LKB1 cKO mice, LKB1 expression was significantly reduced in the cerebellum following Cre recombinase expression (Fig. 1B, Fig. S1B). In contrast, AMPKα1 expression remained largely unchanged, whereas the active zone protein ELKS, which may be regulated by SAD-B, a downstream effector of LKB1 (Barnes et al., 2007; Inoue et al., 2006; Mochida et al., 2016; Shelly and Poo, 2011), showed a slight but statistically insignificant decrease (Fig. 1B, Fig. S1B). Since LKB1 phosphorylates key proteins, thereby regulating metabolic status, it is unlikely to directly affect the expression of downstream factors. Notably, LKB1 cKO mice exhibited reduced survival after weaning, with survival declining to about 50% by 10 weeks and 20% by 20 weeks (Fig. 1C). Similarly, although body weights at weaning were comparable between LKB1 cKO and control mice, weight gain in LKB1 cKO mice plateaued thereafter and remained significantly lower than that of control mice (Fig. 1D). Assessments of activity, food intake, and water consumption in a subset of LKB1 cKO mice did not reveal any obvious factors contributing to the reduced survival.

Because the cerebellum plays a critical role in motor coordination, we evaluated motor performance in LKB1 cKO mice using the rotarod test. At 4–5 weeks of age, LKB1 cKO mice and control mice did not show any significant differences in rotarod performance. In contrast, by 8–9 weeks of age, LKB1 cKO mice developed progressive motor coordination deficits, showing a significantly shorter latency on the rotarod test than control mice (Fig. 1E). This contrasts with a previous report in mice with LKB1 deletion in GCPs, which showed impaired motor coordination and increased stride length at P30, approximately 4 weeks of age (Men et al., 2015). Furthermore, locomotion and gait were analyzed in freely moving mice on a flat arena by tracking multiple body parts using DeepLabCut (Fig. 1F). Analysis of the total movement distance (activity), time spent in the center zone, and walking speed revealed no significant differences between control and LKB1 cKO mice (Fig. 1G, Table S1). Likewise, fore–hind limb distance, the limb projection area, and nose height during walking were comparable between groups. In contrast, tail height progressively increased in LKB1 cKO mice and became significantly higher with age, raising the possibility that tail position was altered to compensate for impaired balance during walking (Fig. 1H and 1I, Table S1). Notably, these differences were specific to walking, as none of these parameters differed significantly during resting (Table S1).

### LKB1 depletion in GCs led to age-dependent degeneration of the ML

To investigate the morphological basis behind the progressive motor dysfunction in LKB1 cKO mice, we analyzed cerebellar morphology using immunofluorescence staining. The ML was visualized by immunostaining for VGluT1, which labels PF terminals to the PCs. At 4–5 weeks of age, shortly after weaning, motor function was largely preserved and the overall cerebellar morphology of LKB1 cKO mice was largely comparable to that of control mice, although mild excessive foliation was observed in the posterior lobules, particularly lobules VIII–X (Fig. 2A). At 8–11 weeks of age, when motor performance began to decline, no major additional morphological changes were apparent. However, at 20–30 weeks of age, when only approximately 20% of LKB1 cKO mice survived and some animals exhibited an abnormal gait with hindlimb dragging, the cerebellum was markedly reduced in size, while excessive foliation remained evident in lobules VIII–IX (Fig. 2A). Previous studies in which LKB1 was depleted in GCPs reported lobular hyperplasia and enlargement of the cerebellar cortex at 4 weeks of age, indicating an important role of LKB1 in GC differentiation and migration (Men et al., 2015; Ryan et al., 2017). In contrast, our LKB1 cKO mice exhibited excessive foliation predominantly in lobules VIII–X (Fig. 2A). This posterior predominance likely reflects the earlier onset of Cre-mediated LKB1 depletion in the posterior lobules during cerebellar development, which may affect GC differentiation (Fig. S1A). The extent and distribution of abnormal foliation varied among younger LKB1 cKO mice (Fig. S2A), likely reflecting individual differences in the timing and spatial progression of Cre expression and subsequent LKB1 depletion in GCPs. In contrast, mice older than 8 weeks consistently exhibited excessive foliation predominantly in lobule IX. The relative absence of anterior foliation in older mice may suggest that mice with more extensive developmental abnormalities were less likely to survive to later ages.

**Figure 2.**
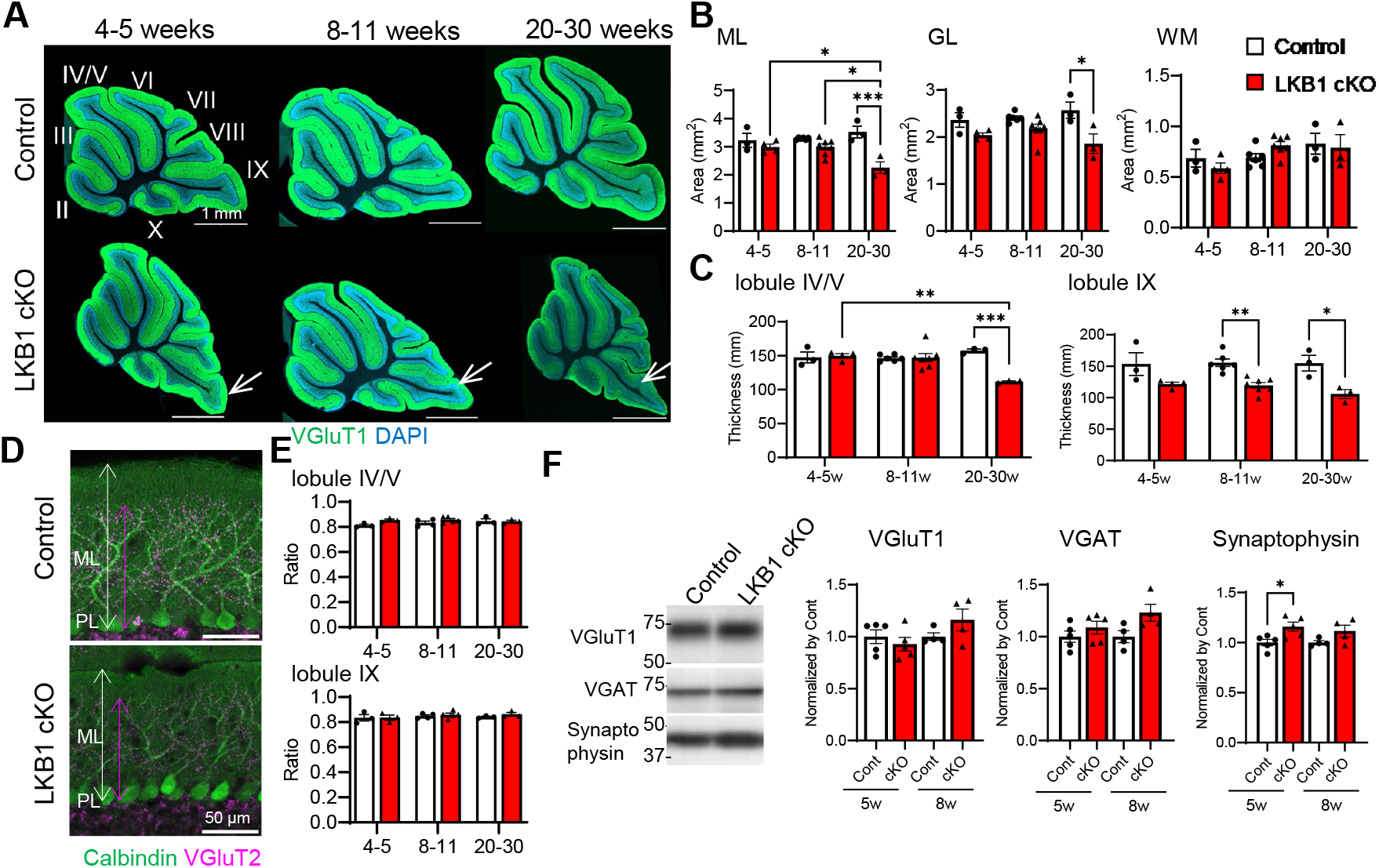
Morphological degeneration of cerebellar ML in the LKB1 cKO mice. (A) Representative cerebellar sagittal sections immunolabeled with anti-VGluT1 antibody (green) and DAPI (blue), shown across development from young (4–5 weeks) to aged (20–30 weeks) control and LKB1 cKO mice. The cerebellum of LKB1 cKO mice exhibited excessive foliation of the posterior lobules (lobule VIII, IX; white arrows). (B-C) Quantification of layer areas showed an age-dependent reduction in the molecular layer (ML) and granule cell layer (GL) in LKB1 cKO mice at 20-30 weeks, while the white matter (WM) remained unchanged. ML thickness was reduced in lobule IV/V of LKB1 cKO mice at 20–30 weeks and in lobule IX at an earlier stage (8-11 weeks), indicating region- and age-specific degeneration. Control; 4–5 weeks = 3, 8–11 weeks = 6, 20–30 weeks = 3, LKB1 cKO; 4–5 weeks = 4, 8–11 weeks = 7, 20–30 weeks = 3. Mean ± SEM, * p < 0.05, ** p < 0.01, *** p < 0.001, two-way ANOVA post-hoc Tukey test. (D-E) Representative images of Purkinje cells labeled with anti-calbindin antibody (green) and climbing fibers labeled with anti-VGluT2 antibody (magenta). The relative extent of climbing fiber innervation to Purkinje cell dendrites was not significantly different between the control and LKB1 cKO. Control; 4–5 weeks = 3, 8–11 weeks = 4, 20–30 weeks = 3, LKB1 cKO; 4–5 weeks = 3, 8–11 weeks = 5, 20–30 weeks = 3. Mean ± SEM, two-way ANOVA post-hoc Tukey test. (F) Immunoblotting of cerebellar homogenates revealed no marked changes in the synaptic protein expression, including VGluT1 (excitatory terminals), VGAT (inhibitory terminals), and actin. At younger age, the expression of synaptophysin (general synaptic marker) was slightly increased in LKB1 cKO mice. 5 weeks; Control = 5, LKB1 cKO = 5, 8 weeks; Control = 4, LKB1 cKO = 4. Mean ± SEM, * p < 0.05, Student’s t-test.

Quantitative analysis showed that the total areas of the ML, GL, and white matter (WM) in sagittal sections of the cerebellar vermis did not differ significantly between control and LKB1 cKO mice at 4–5 or 8–11 weeks of age. However, the areas of the ML and GL were significantly reduced in LKB1 cKO mice after 20 weeks of age (Fig. 2B). Furthermore, although ML thickness in the anterior lobules (lobules IV/V) was significantly reduced only after 20 weeks, ML thinning in the posterior lobule (lobule IX) was already evident at 8–11 weeks of age (Fig. 2B). These findings suggest that cerebellar shrinkage began in the posterior lobules, where Cre-mediated recombination occurred earlier, and progressively extended to the anterior cerebellum with age. In addition, cell density in the ML was significantly higher in LKB1 cKO mice in both the anterior and posterior lobules (Fig. S2B-C). This increased cell density may reflect impaired migration of GCPs caused by developmental loss of LKB1, consistent with previous reports demonstrating a role for LKB1 in GCP migration (Men et al. 2015; Ryan et al. 2017).

In the ML, PCs also receive CF inputs, which originate from the inferior olive and innervate the proximal dendritic spines of PCs. Immunofluorescence staining with an anti-calbindin antibody to label PCs and an anti-VGluT2 antibody to visualize CF terminals revealed the relative extent of CF innervation along PC dendrites was not significantly altered (Fig. 2D-E, Fig. S2D). These results indicate that LKB1 deficiency in GCs predominantly affected PF–PC synapses, with no detectable alteration in CF–PC innervation. Furthermore, analysis of synaptic protein expression in cerebellar homogenates showed that synaptophysin was slightly increased in LKB1 cKO mice at 5 weeks of age, whereas VGluT1, a marker of PF terminals, and VGAT, a marker of inhibitory terminals, were not significantly altered (Fig. 2F).

### PF–PC synaptic alterations precede ML atrophy following LKB1 loss in GCs

Microglia were traditionally regarded as immune cells responsible for the phagocytic clearance of damaged cells, and recent studies have revealed their complex and multifunctional roles in synaptic remodeling under both physiological and pathological conditions (Iida, Tanaka, and Okabe 2019; Pereira-Iglesias et al. 2025). Immunofluorescence staining for microglia revealed a significant increase in microglial density in the ML, but not in the GL, of LKB1 cKO mice (Fig. 3A-C). In lobules IV/V, microglial density in the ML progressively increased in LKB1 cKO mice and was significantly higher than that in control mice from 8 weeks of age onward (Fig. 3B). In contrast, in lobule IX, microglial density in the ML was already increased at 4–5 weeks of age (Fig. 3B). Despite these changes, total Iba1 expression in cerebellar homogenates showed a slight but statistically nonsignificant increase in LKB1 cKO mice compared with controls (Fig. 3D). These findings suggest that LKB1 depletion in GCs induced a regionally progressive microglial response in the ML, potentially associated with alterations in PF–PC synapses.

**Figure 3.**
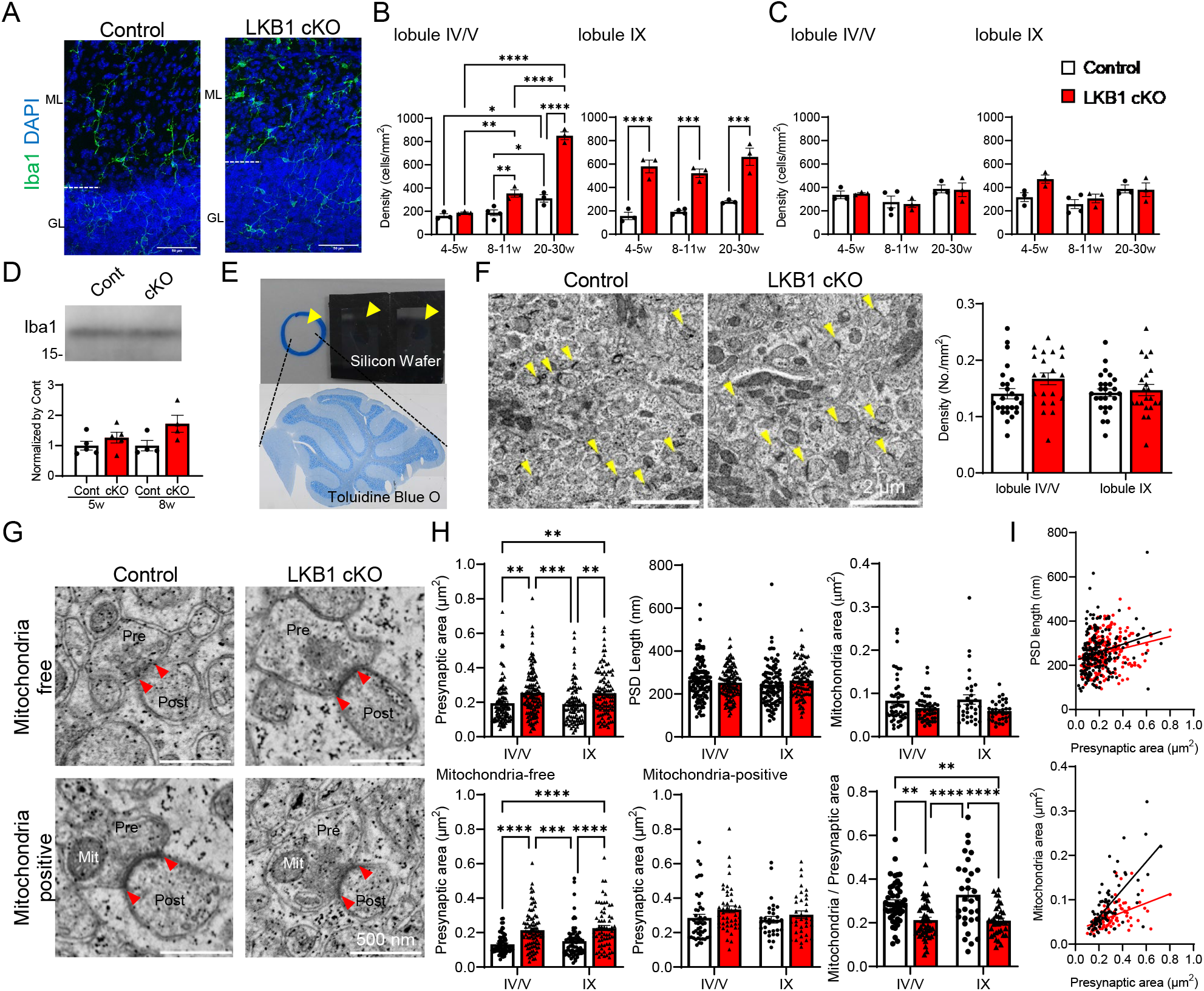
Loss of LKB1 in GCs increased microglial density and induced ultrastructural changes characterized by enlargement of MT-free PF–PC synapses. (A) Representative images of cerebellar sections immunolabeled with anti-Iba1 antibody showed increased microglial density in LKB1 cKO mice. (B-C) Microglial density in the molecular layer (ML) indicated an age-dependent increase in the anterior lobule (IV/V) of LKB1 cKO mice. In the posterior lobule (IX), the density was already elevated at a younger age in LKB1 cKO mice. Microglial density in the granule cell layer (GL) was not altered. Control; 4–5 weeks = 3, 8–11 weeks = 4, 20–30 weeks = 3, LKB1 cKO; 4–5 weeks = 3, 8–11 weeks = 3, 20–30 weeks = 3. Mean ± SEM, ** p < 0.01, *** p < 0.001, **** p < 0.0001, two-way ANOVA post-hoc Tukey test. (D) Immunoblotting of cerebellar homogenate showed a slight but not significant increase in Iba1 expression in LKB1 cKO mice. Control; 5 weeks = 5, 8 weeks = 4, LKB1 cKO; 5 weeks = 5, 8 weeks = 4. Mean ± SEM, Student’s t-test. (E) To quantify the fine structure of PF-PC synapses in lobule IV/V and lobule IX using section-SEM technique, semi-thin sagittal sections of the whole cerebellum were collected on glass slides or silicon wafers (yellow arrowheads). Sections collected on glass slides were stained with toluidine blue O to visualize the overall morphology of the cerebellum under a light microscope. (F) Representative section-SEM images at the ML showed numerous PF-PC synapses, identified by the accumulation of synaptic vesicles adjacent to a postsynaptic PSD (yellow arrowheads). The PF-PC synapse density was not significantly different between groups. Control; n = 25 images from 3 mice, LKB1 cKO; n = 21 images from 3 mice. Mean ± SEM, two-way ANOVA post-hoc Tukey test. (G) Detailed analysis of PF-PC synapses, including presynaptic area, PSD length (distance between two red arrowheads), and presynaptic MT area. (H) The presynaptic area was increased in the LKB1 cKO mice in both lobule IV/V and IX, while PSD length and MT area were not significantly different. When presynaptic terminals were categorized based on the presence of MT, the presynaptic area was increased in MT-free terminals in LKB1 cKO mice. Although the size of MT-positive terminals was unchanged, the proportion of MT area within terminals was significantly decreased in LKB1 cKO mice. n = 100–130 synapses from Control; n = 3 mice, LKB1 cKO; n = 3 mice. Mean ± SEM, ** p < 0.01, *** p < 0.001, **** p < 0.0001, two-way ANOVA post-hoc Tukey test. (I) Correlation between presynaptic area and PSD length was similar in control and LKB1 cKO mice (Pearson’s R; Control = 0.23, LKB1 cKO = 0.23). However, the correlation between the presynaptic and MT areas was reduced in the LKB1 cKO mice (Pearson’s R; Control = 0.69, LKB1 cKO = 0.47).

To further investigate the effects of LKB1 loss in GCs on PF–PC synapses, we examined the ultrastructure of the ML. Semithin sagittal sections of the whole cerebellum were used to identify the regions of interest (the ML in lobules IV/V and IX), for subsequent observation by section scanning electron microscopy (section-SEM) using backscattered electron imaging (Fig. 3E). PF–PC synapses were identified by presynaptic terminals containing clusters of synaptic vesicles adjacent to the active zone and contacting a single postsynaptic spine containing a postsynaptic density (PSD). The density of PF–PC synapses was not significantly different between control and LKB1 cKO mice (Fig. 3F). At higher magnification (×40,000), the mean PSD length was comparable, whereas the area of presynaptic terminals was enlarged in LKB1 cKO mice (Fig. 3G-H). Similar alterations were observed in both the anterior and posterior lobules.

LKB1 phosphorylates and activates multiple downstream kinases, some of which contribute to mitochondrial biogenesis in the presynaptic terminal (Courchet et al. 2013; Kwon et al. 2016). Since mitochondria (MT) play an important role in maintaining synaptic function and neurotransmitter release, we further examined MT within PF terminals. Although the MT area within individual PF terminals and the size of MT-containing presynaptic terminals was comparable between control and LKB1 cKO mice, presynaptic terminals lacking MT were significantly larger in LKB1 cKO mice (Fig. 3G-H). Moreover, the proportion of the presynaptic terminal area occupied by MT was significantly lower in LKB1 cKO mice (Fig. 3G and 3H). Consistent with these findings, the correlation between the presynaptic terminal area and MT area was lower in LKB1 cKO mice, while the correlation between the presynaptic terminal area and PSD length was comparable (Fig. 3I). Together, these ultrastructural changes indicated that LKB1 depletion in GCs disrupted the structural organization of PF presynaptic terminals, particularly the relationship between terminal size and MT content.

### 3D reconstruction of PFs revealed that loss of LKB1 in GCs increased multisynaptic contacts on the PF boutons

Conventional section-SEM analysis provides quantitative information on a large number of synapses based on two-dimensional (2D) images; however, three-dimensional (3D) analysis is required to understand the overall organization of PF synapses. In LKB1 cKO mice, degeneration was more pronounced in the posterior lobule IX, whereas the ultrastructural features of individual synapses were largely comparable between lobules IV/V and IX. Therefore, serial block-face SEM (SBF-SEM) was performed on the ML of lobule IX. First, typical PF–PC synapses were identified in serial SEM images, and the corresponding PFs were traced through consecutive sections (500–600 images) for 3D reconstruction (Fig. 4A-B). Given a section thickness of 50 nm, the reconstructed PF length was approximately 25–30 μm along the z-axis. Using a cross-sectional area >0.15 μm^2^ with at least five consecutive sections as the criteria for PF bouton (BT) identification, we analyzed 84 BTs from 21 PFs in control mice and 79 BTs from 19 PFs in LKB1 cKO mice (Fig. 4C). The volume and surface area of the reconstructed BTs and the MT contained within them were not significantly different between control and LKB1 cKO mice (Fig. 4C). Interestingly, 20–30% of BTs lacked MT in both control and LKB1 cKO mice, indicating that this feature was not specific to LKB1 depletion. However, the MT volume fraction within individual BTs was significantly higher in LKB1 cKO mice (Fig. 4C). Because some BTs contained no MT, whereas others contained multiple MT, we calculated the MT/BT volume ratio for individual MT, both including and excluding BTs without MT (Fig. S3A). The MT/BT volume ratio was also calculated using the summed MT volume within each BT, with or without including BTs lacking MT. In all cases, the MT/BT volume ratio was significantly higher in LKB1 cKO mice (Fig. S3A). In contrast, the distances between adjacent BTs were not significantly different between control and LKB1 cKO mice, consistent with the 2D analysis showing no significant difference in PF–PC synapse density between genotypes.

**Figure 4.**
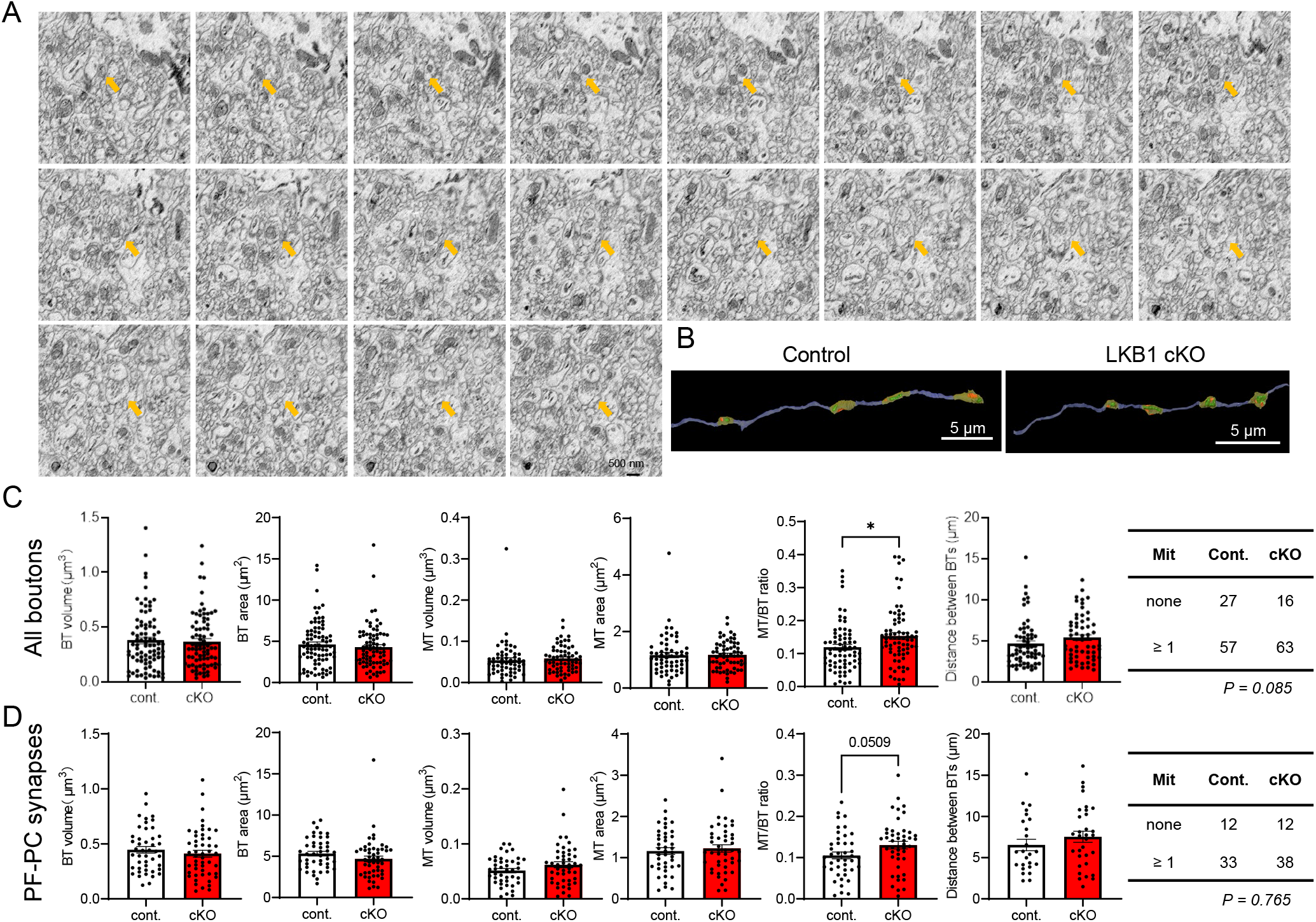
Loss of LKB1 in GCs increased MT occupancy in PF boutons revealed by 3D SBF-SEM analysis. (A-B) Representative serial SBF-SEM images of the cerebellar ML acquired at 50-nm section thickness. Orange arrows indicate a representative PF–PC synapse and the corresponding PF traced through consecutive sections. 3D reconstruction of the traced PF revealed multiple BTs, which were defined as PF swellings with a cross-sectional area ≥ 0.15 μm^2^ that persisted for at least five consecutive sections. (C) Quantitative analysis of PF BT and MT morphology, including BT volume and surface area, MT volume and surface area, the MT/BT volume ratio, and the distance between adjacent BTs as an indicator of BT density along PFs. The table summarizes the proportions of BTs with or without MT, which were compared between Control and LKB1 cKO mice using the chi-square test. Control; n = 84 BTs from 21 PFs, 2 ROI from 2 mice, cKO; n = 79 BTs from 19 PFs, 2 ROI from 2 mice, Mean ± SEM, * p < 0.05, Student’s t-test. (D) Similar morphological analyses were performed specifically for PF boutons forming PF–PC synapses, which were identified by synaptic contacts with PC spines. BT volume and surface area, MT volume and surface area, and the MT/BT volume ratio were quantified and compared between Control and LKB1 cKO mice. Control; n = 45 BTs from 20 PFs, cKO; n = 50 BTs from 19 PFs, Mean ± SEM, * p < 0.05, Student’s t-test.

The reconstructed BTs were classified into four categories according to their postsynaptic targets and synaptic connectivity: BTs forming synapses with PC spines (PF–PC synapses), BTs forming synapses with inhibitory dendrites, BTs forming synapses with both PC spines and inhibitory dendrites, and BTs without an identifiable synaptic connection (Fig. 4D, Fig. S3C). These four categories were subsequently analyzed separately. The majority of BTs formed PF–PC synapses, with no significant differences in BT or MT volume and surface area between genotypes. The MT volume ratio was slightly higher but not significantly different (Fig. 4D). The distances between PF–PC synaptic BTs were also not significantly different between genotypes. Similarly, among BTs forming synapses with inhibitory dendrites and those forming synapses with both PC spines and inhibitory dendrites, none of the examined BT or MT parameters differed significantly between control and LKB1 cKO mice (Fig. S3C). Although conventional 2D SEM analysis revealed an enlargement of PF terminals lacking MT, 3D reconstruction showed that the proportion of BTs without MT was comparable between control and LKB1 cKO mice (Fig. 4D; chi-square test, p = 0.76). Further 3D analysis revealed a slight but not significant reduction in the Z-axis elongation of MT in LKB1 cKO mice (Fig. S3B). In addition, analysis of the MT distribution within BTs based on centroid distances showed that MT were located closer to the center of BTs in LKB1 cKO mice (Fig. S3D). These morphological changes may alter the spatial relationship between MT and active zones, potentially explaining the enlarged PF terminals lacking MT in section-SEM. Therefore, quantitative 3D analysis indicated a limited role for LKB1 in regulating presynaptic MT morphology. In contrast, 3D reconstruction revealed a significant increase in the number of active zones per BT in LKB1 cKO mice, particularly among BTs forming PF–PC synapses (Fig. 5A-B, Fig. S4). The ratio of BTs forming synaptic contacts with PC spines to those contacting inhibitory dendrites increased from 2.8:1 in control mice to 6.25:1 in LKB1 cKO mice, although this difference was not statistically significant (chi-square test, p = 0.091; Fig. S4A). This trend suggests a possible shift toward preferential synaptic connectivity with PC spines following LKB1 deletion. BTs are generally considered to form a single synaptic contact, and occasionally two, with PC spines (Ichikawa, Sakimura, and Watanabe 2016; Loschky et al. 2022), although a recent 3D SEM study demonstrated more complex PF–PC connectivity, including multiple synaptic contacts arising from individual PFs (Loschky et al. 2022). Consistent with alterations in this connectivity, we found that some PF BTs in LKB1 cKO mice formed multiple synaptic contacts with three or four PC spines (Fig. 5B). We further examined the relationship between the BT size and the active zone area. The BT surface area was positively correlated with the active zone area in control mice, whereas they were negatively correlated in LKB1 cKO mice (Fig. 5C). These findings indicate that LKB1 loss increased the number of synaptic contacts formed by individual BTs and altered the relationship between BT size and synaptic area, suggesting altered PF–PC connectivity that may influence PC activity and cerebellar output.

**Figure 5.**
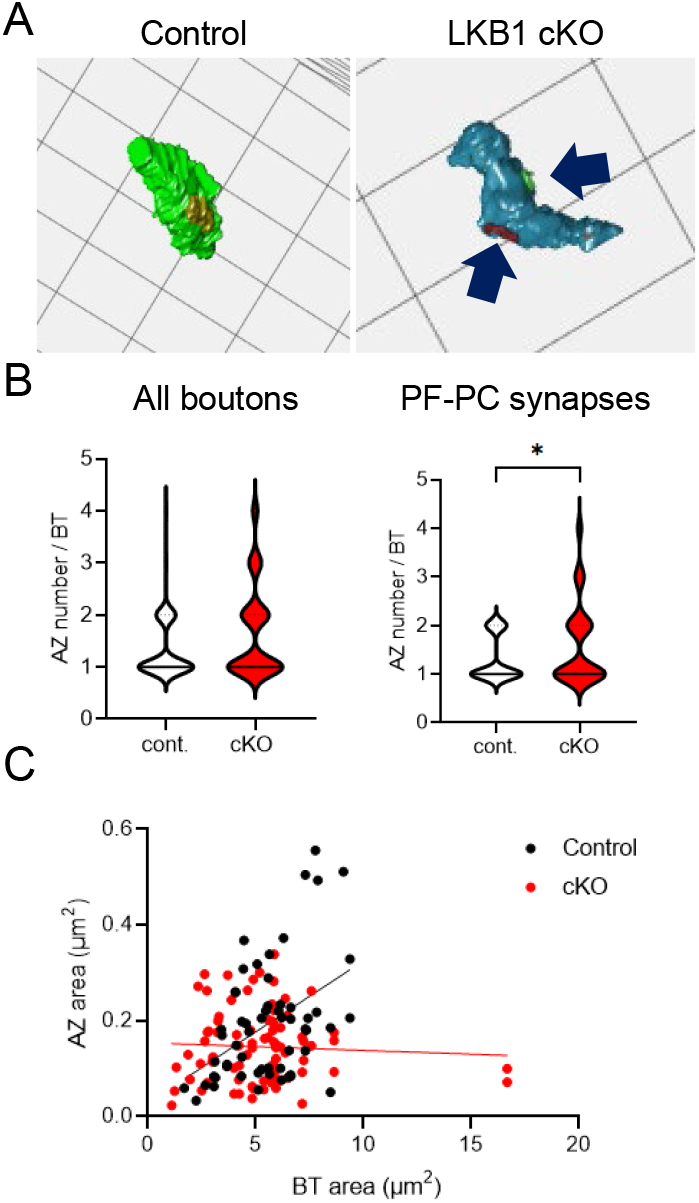
Loss of LKB1 in GCs increased multisynaptic PF boutons at PF–PC synapses. (A) Representative 3D reconstructed BTs with a single active zone in control mice and two active zones in LKB1 cKO mice. Active zones were identified by the accumulation of presynaptic vesicles adjacent to the presynaptic membrane and the presence of a corresponding postsynaptic density (PSD) at the apposed PC spine membrane. (B) Quantitative analysis showed that the number of active zones per BT was significantly increased in LKB1 cKO mice in PF–PC synapses. Mean ± SEM, * p < 0.05, Student’s t-test. (C) Correlation analysis revealed a reduction in the strength of the relationship between BT size and active zone area in LKB1 cKO mice.

### Cerebellar dysfunction was associated with muscle deterioration and reduced neuromuscular innervation

Cerebellar dysfunction as seen in *Car8^wdl^* mice and GluD2 or Cbln1 KO mice exhibiting cerebellar dysfunction with abnormal neural activity in PCs causes impaired contraction in skeletal muscles (Matsuda et al. 2010; Miterko et al. 2019, 2021; Takeuchi et al. 2018). To investigate whether progressive thinning of cerebellar ML and altered PF–PC synaptic connectivity were associated with changes in skeletal muscles, we analyzed hindlimb muscles in LKB1 cKO mice. The weights of hindlimb skeletal muscles, including the tibialis anterior (TA), extensor digitorum longus, and gastrocnemius and soleus normalized to body weight were not significantly different between control and LKB1 cKO mice, regardless of age or sex (Fig. 6A-B). However, the cross-sectional area of TA muscle fibers at 8–10 weeks of age, when motor coordination deficits became evident, was significantly reduced in male LKB1 cKO mice, whereas no significant difference was observed in female mice (Fig. 6C-D). Furthermore, muscle fiber density was significantly increased in both male and female LKB1 cKO mice (Fig. 6E). These results suggest that cerebellar dysfunction in LKB1 cKO mice was associated with alterations in skeletal muscle morphology, including reduced muscle fiber size, coinciding with the onset of motor coordination impairments.

**Figure 6.**
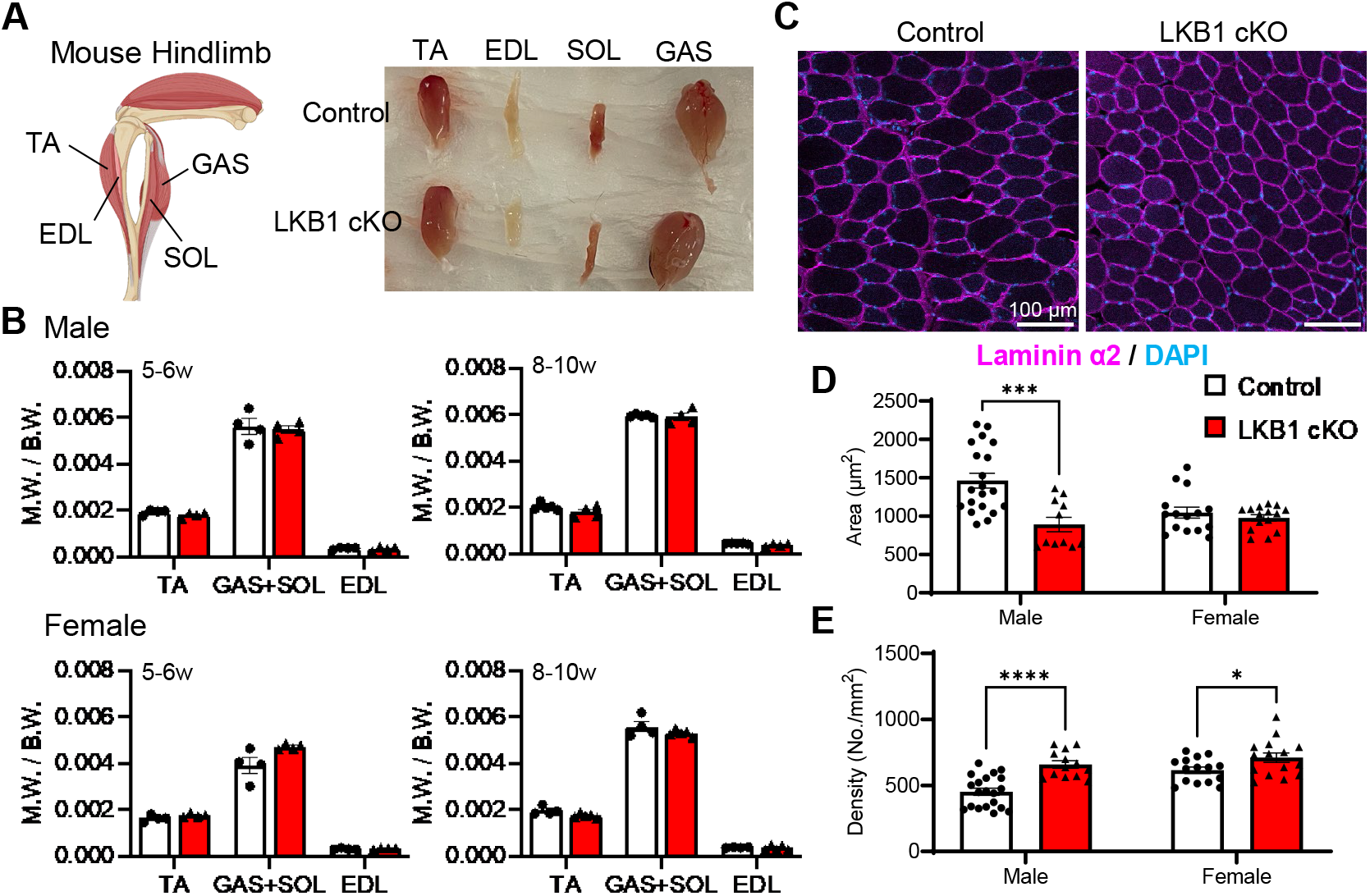
Evaluation of muscle atrophy in the skeletal muscles of LKB1 cKO mice. (A) Schematic diagram of hindlimb skeletal muscles and harvested samples from 8-week control and LKB1 cKO mice. (B) Muscle weight normalized to body weight showed no significant difference. Control; 5–6 weeks = 8 (male 4, female 4), 8–10 weeks = 9 (male 5, female 4), LKB1 cKO; 5–6 weeks = 8 (male 4, female 4), 8–10 weeks = 9 (male 4, female 5), Mean ± SEM, Student’s t-test. (C) Representative images of cross-sectioned muscle fibers immunolabeled with anti-laminin α2 antibody (magenta) with DAPI. Cross-sectional area was reduced, and the density was increased in LKB1 cKO mice, with the change being pronounced in males. Control = 7 (male 4, female 3), LKB1 cKO = 6 (male 3, female 3), 8–10 weeks, Mean ± SEM, * p < 0.05, *** p < 0.001, **** p < 0.0001, Student’s t-test.

Given the close association between muscle atrophy and NMJ integrity, we examined NMJ morphology by labeling postsynaptic acetylcholine receptors with α-bungarotoxin and presynaptic terminals with an anti-synaptophysin antibody (Fig. 7A). The size (length and width) and area of the postsynaptic site were not significantly different in LKB1 cKO mice and control mice (Fig. 7B). However, the presynaptic terminal area and the overlap between presynaptic and postsynaptic sites were significantly reduced in 8–10-week-old LKB1 cKO mice, leading to a marked reduction of presynaptic innervation at the NMJs (Fig. 7C-D). Furthermore, the proportion of NMJs with an innervation ratio below 10% was comparable between genotypes at 5–6 weeks but increased to over 15% in LKB1 cKO mice by 8–10 weeks (Fig. S5A). Despite these alterations in NMJ innervation, grip strength at 8–10 weeks of age was not significantly different between control and LKB1 cKO mice (Table S2). In an attempt to determine the contribution of LKB1 depletion in motor neurons, western blotting and immunohistochemistry revealed no detectable Cre expression in the spinal cord, in contrast to robust Cre expression in cerebellar GCs (Fig. S5B-C). Additionally, neither the density of motor neurons in the L3 spinal cord nor the number of retrogradely labeled motor neurons innervating the TA differed between the LKB1 cKO mice and controls (Fig. S5D-E). Together, these findings suggest that cerebellar dysfunction in LKB1 cKO mice was accompanied by alterations in NMJ organization, which may contribute to the observed motor impairments.

**Figure 7.**
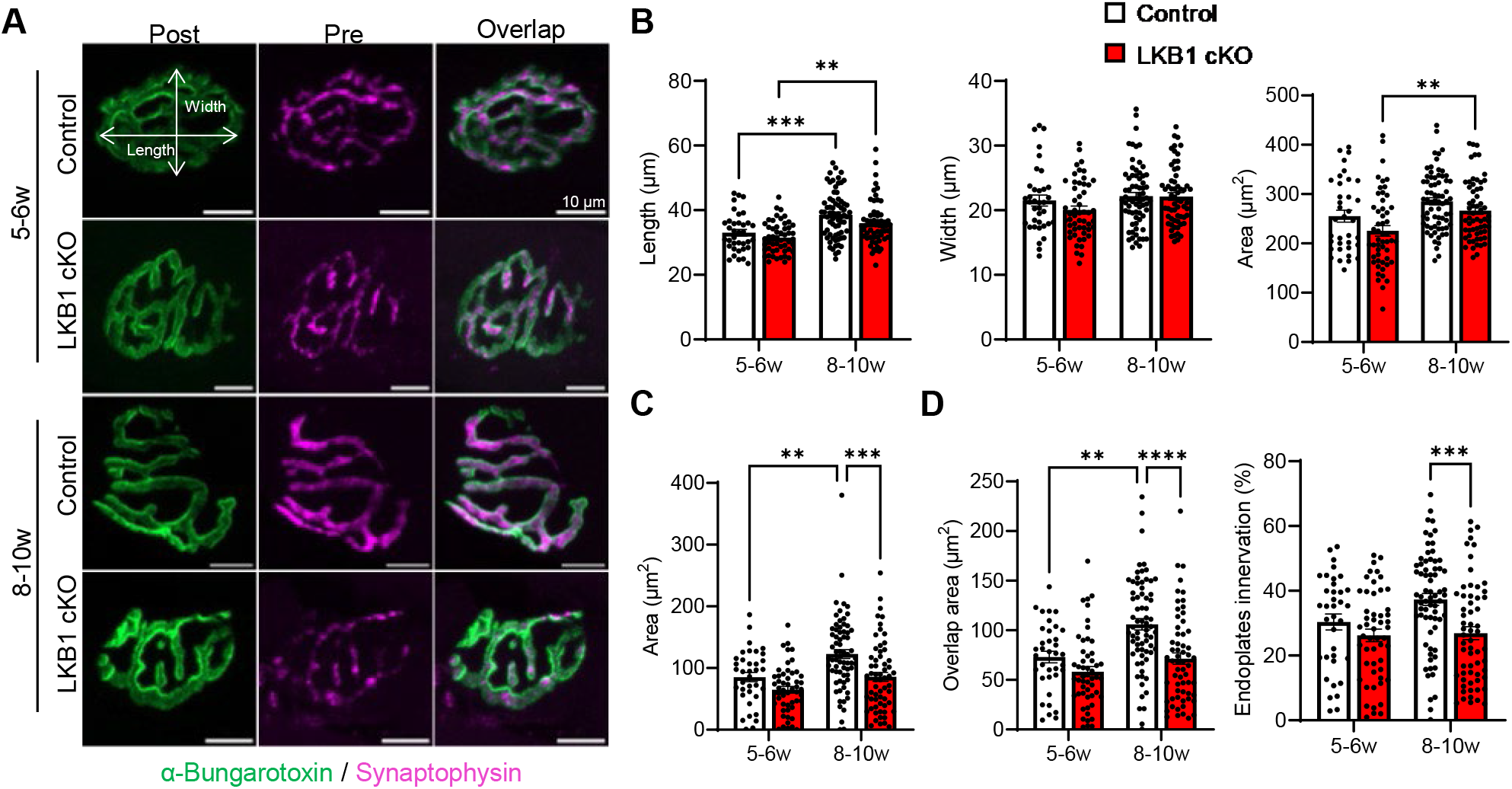
Reduced presynaptic innervation at neuromuscular junctions in LKB1 cKO mice. (A) Representative images of neuromuscular junctions in TA muscles. Postsynaptic sites were labeled with α-bungarotoxin (green), and presynaptic terminals were labeled with anti-synaptophysin antibody (magenta). (B) Quantification of postsynaptic terminal size, measured as length (long axis), width (short axis), and area. The postsynaptic terminal length was increased with age in both control and LKB1 cKO mice. (C) Measurement of the presynaptic terminal area revealed a slight age-dependent increase in control mice, whereas in LKB1 cKO mice, the area remained unchanged and was significantly decreased at 8–10 weeks of age. n = 36–69 neuromuscular junctions from control; 5–6 weeks = 3, 8–10 weeks = 6 mice, LKB1 cKO; 5–6 weeks = 5, 8–10 weeks = 5 mice, Mean ± SEM, ** p < 0.01, *** p < 0.001, **** p < 0.0001, Student’s t-test.

## DISCUSSION

The cerebellum plays essential roles in coordinating movements by correcting errors between intended motor commands and actual movement, thereby ensuring smooth execution of motor sequences. Cerebellar dysfunction leads to various motor deficits, including gait abnormalities and asymmetric limb positioning (Jaarsma et al. 2024; Machado et al. 2015, 2020; Takeuchi et al. 2018). Studies have shown that embryonic deletion of LKB1 from GCPs caused motor deficits as early as 4 weeks of age (Men et al. 2015; Ryan et al. 2017). In contrast, the present study, which involved postnatal depletion of LKB1, revealed no significant motor abnormalities at 4–5 weeks. However, impaired motor coordination became evident at 10 weeks of age, indicating a progressive deterioration of motor function. Notably, no temporal changes in limb positioning during gait were observed, suggesting that the coordination deficits were not due to overt gait disturbances (Fig. 1).

During cerebellar development, LKB1 plays multiple critical roles, including promoting GC proliferation and migration, which causes cerebellar cortical hypertrophy, as well as regulating dendritic self-avoidance in PCs (Kuwako and Okano 2018a; Men et al. 2015; Ryan et al. 2017). In the present postnatal LKB1 depletion in GCs, excessive foliation was observed primarily in lobule IX (Fig. 2), likely because Cre recombination driven by E3CreN mice began in posterior lobules (Fig. S1). Moreover, LKB1 deletion in GCs increased the proportion of PF BTs forming multiple synaptic contacts, particularly those containing three or more active zones, while reducing the correlation between the active zone area and the BT surface area (Figs. 4-5). Together with the previously reported role of LKB1 in dendritic self-avoidance in PCs (Kuwako and Okano 2018a), these findings suggest that LKB1 may also contribute to the proper organization and restriction of synaptic contacts at presynaptic PF BTs. Although the structural and functional properties of multisynaptic BTs at PF–PC synapses have not been extensively investigated, increased multisynaptic BT has been reported in tottering mice carrying mutations in P/Q-type Ca^2+^ channels (Kim et al. 2019), along with impaired PF–PC synaptic transmission (Matsushita et al. 2002). Thus, the structural alterations observed in LKB1 cKO mice may affect PF–PC synaptic transmission and contribute to impaired motor coordination.

### Age-dependent cerebellar degeneration caused by less presynaptic signal transduction of LKB1

Postnatal depletion of LKB1 from GCs led to progressive degeneration of ML in cerebellum, initially appearing in the posterior lobules at around 8–10 weeks of age and gradually spreading to anterior lobules by 20–30 weeks, indicating age-dependent cerebellar degeneration (Fig. 2). Notably, an increase in microglial density—an indicator of neuroinflammation—was observed prior to ML thinning (Fig. 3), suggesting that microglia may respond to early synaptic abnormalities and contribute to degenerative processes.

Synaptic neurotransmission is a highly energy demanding process, requiring localized ATP production and calcium buffering, functions typically supported by presynaptic MT (Chavan et al. 2015; Kwon et al. 2016). One key pathway, the LKB1–NUAK1 kinase pathway, has been shown to regulate axon branching by promoting MT immobilization at terminals (Courchet et al. 2013). Conventional 2D SEM analysis revealed a reduced correlation between the presynaptic area and MT area in LKB1 cKO mice (Fig. 3), suggesting alterations in presynaptic MT organization. However, 3D reconstruction of EM revealed no significant differences in BT volume, MT volume, or the proportion of BTs lacking MT, which accounted for approximately 25% in control and LKB1 cKO mice (Fig. 4). The discrepancy between 2D and 3D analyses may be partly explained by differences in MT geometry and the spatial arrangement of active zones. In particular, the increased prevalence of multisynaptic BTs in LKB1 cKO mice may influence the cross-sectional profiles obtained from sections containing active zones. Thus, the changes observed in 2D EM may reflect alterations in synaptic organization rather than an overall enlargement of presynaptic BTs. Although LKB1 deletion in GCs had limited effects on presynaptic MT morphology, functional alterations cannot be excluded, as LKB1 has been reported to regulate the expression of the mitochondrial calcium uniporter (MCU) (Kwon et al. 2016). Thus, LKB1 deficiency may impair presynaptic calcium homeostasis, potentially affecting glutamate release at PF–PC synapses and subsequent PC activity. These alterations may contribute to microglial activation and neuroinflammation, ultimately leading to cerebellar degeneration and motor deficits.

### Pathophysiological mechanisms for cerebellar ataxia in mouse models

We applied deep learning-based methods for gait analysis on recordings of freely moving mice from below using an RGB-D camera (Oikawa et al. 2023; Tsuruda et al. 2023). Using 3D tracking of key body points, including the limbs, nose, and tail, we found that LKB1 cKO mice showed no significant alterations in gait parameters except for a higher tail position (Fig. 1). Previous studies using cerebellar mutant mice have demonstrated alterations in tail and nose movements during locomotion. For example, PC degeneration model mice exhibit exaggerated tail and nose oscillations, whereas reeler model mice show impaired gait coordination associated with disrupted cerebellar circuitry (Darmohray et al. 2019; Machado et al. 2015, 2020). In contrast, SCA6 transgenic mice exhibit reduced tail and nose elevation during balance beam walking (Piotrowski et al. 2024). Therefore, the changes in tail hight in LKB1 cKO mice may be associated with disrupted PF–PC synaptic organization, potentially affecting cerebellar output and motor coordination. Furthermore, disruption of PF–PC synaptic integrity can contribute to cerebellar dysfunction. Mouse models lacking GluD2 in PCs or Cbln1 in GCs demonstrate the importance of PF–PC synapses in motor coordination, as these mice exhibit reduced PF–PC synapse density and impaired synaptic plasticity, including long-term depression (Hirai et al. 2005; Ichikawa et al. 2016; Kakegawa et al. 2018; Yuzaki 2013). Since Cbln1 forms a trans-synaptic complex with presynaptic neurexins and postsynaptic GluD2, in adult Cbln1-deficient mice, cerebellar administration of recombinant Cbln1 has been shown to restore PF–PC synapses and improve motor function (Matsuda et al. 2010; Suzuki et al. 2020; Yuzaki 2010).

### Limitations of the study

In this study, LKB1 depletion in cerebellar GCs was associated with presynaptic abnormalities at PF–PC synapses, and, unexpectedly, at NMJs, with the latter preceding skeletal muscle atrophy (Figs. 6-7, Fig. S5). This temporal sequence is consistent with previous studies reporting that NMJ degeneration can precede muscle atrophy (Liu et al. 2017; Rodríguez Cruz et al. 2020). Although we confirmed that Cre expression was not detected in spinal motor neurons, potential LKB1 depletion in other brain regions was not examined. Furthermore, while cerebellar output through the red nucleus and reticulospinal pathways is known to modulate spinal motor neuron activity, it remains unclear whether alterations in PF–PC synaptic organization in LKB1 cKO mice affect cerebellar output and downstream motor pathways, thereby contributing to the observed peripheral neuromuscular abnormalities.

### Reduced survival and potential seizure-related mortality in LKB1 cKO mice

LKB1 cKO mice exhibited a marked reduction in survival, with over 50% mortality observed by 9–10 weeks of age (Fig. 1). Our preliminary observation using retrospective monitoring of food and water intake as well as activity levels showed less prominent changes, suggesting that lethality was not due to progressive wasting. Notably, some LKB1 cKO mice displayed seizure-like behaviors, raising the possibility that cerebellar dysfunction may have contributed to sudden seizure-related mortality. In previous studies, ducky mice carrying mutations in *Cacna2d2* exhibited absence-like seizures and ataxia due to impaired PC calcium signaling (Barclay et al. 2001). Similarly, stargazer, tottering, and lethargic mutant mice with disrupted calcium channel subunits showed both cerebellar ataxia and spike-wave discharges, often associated with PC dysfunction (Fletcher and Frankel 1999; Schirmer et al. 2022). In the present LKB1 cKO mice, altered PC activity due to synaptic dysregulation may have led to abnormal cerebellar output and increased seizure risk. However, in E3CreN mice, Cre recombination is driven beyond the cerebellum, including in subsets of cortical, hippocampal, thalamic, and brainstem neurons, and contributions of this to the phenotype remain possible (Miyazaki et al. 2012). Additionally, LKB1 effectors such as SIKs have been linked to cortical epileptogenesis (Huang and Li 2022). Therefore, both cerebellar and broader brain circuit dysfunction may underlie the reduced survival and potential seizure-related mortality in LKB1 cKO mice. In summary, our study reveals that LKB1 depletion in cerebellar GCls affected remodeling of PF–PC synaptic organization, progressive cerebellar degeneration, and impaired motor coordination. These findings provide new insights into the role of LKB1 in maintaining cerebellar synaptic architecture and function.

## RESOURCE AVAILABILITY

### Lead contact

Requests for further information and resources should be directed to and will be fulfilled by the lead contact, Akari Hagiwara.

### Materials availability

This study did not generate new unique reagents. Materials used in this study are available from the lead contact upon reasonable request.

### Data and code availability

All source data for gait analysis can be found in Table S1. This paper does not report original code. Any additional information required to reanalyze the data reported in this paper is available from the lead contact upon request.

## ACKNOWLEDGMENTS

This work was supported by JSPS KAKENHI grant 23K23980 to T. Ohtuska, JP22H04926 for advanced bioimaging support (ABiS), and Life Science Foundation of Japan, Soikinome Project of Tokyo University of Science to A. Hagiwara. We wish to thank all the members of the Hagiwara and Ohtsuka laboratories for their helpful discussions and technical assistance, and Dr. Y. Mizunoe and Dr. S. Yanagida (Tokyo University of Science) for expert technical assistance of muscle experiments, Dr. M. Abe and K. Sakimura (Niigata University) for the E3CreN mice, and A. Imai, N. Hattori and H. Ishihara (National Institute for Physiological Sciences) for technical support.

## AUTHOR CONTRIBUTIONS

Conceptualization: R.K., S.S., A.H.; Methodology: R.K., S.S., H.K., S.H., M.K., M.Y., M.W., M.S., K.T., N.O., H.T., A.H.; Investigation: R.K., S.S., H.K., S.H., M.K., M.W., M.S., K.T., N.O., A.H.; Software: M.Y., H.T.; Formal Analysis: R.K., S.S., H.K., N.O., A.H.; Visualization: R.K., S.S., H.K., N.O., A.H.; Writing – Original Draft: R.K., S.S., A.H.; Writing – Review & Editing: A.H., T.O.; Supervision: A.H., T.O.

## DECLARATION OF INTERESTS

The authors declare no competing interests.

## SUPPLEMENTAL INFORMATION

Document S1. Supplemental Figures S1–S5 Table S1. Gait analysis data

Table S2. Grip strength data

## METHODS

### Animals

The Institutional Committee for the Care and Use of Experimental Animals at the Tokyo University of Science (protocol # 1991 and # N22013) and the University of Yamanashi (protocol # A30-21) approved the use of animals. All experiments were conducted according to the recommendations in the Guidelines for Proper Conduct of Animal Experiments of the Science Council of Japan. All efforts were made to reduce the number and suffering of animals used. To generate LKB1 cKO mice, LKB1-flox mice (The Jackson Lab, #014143) (Nakada et al. 2010) were crossed with E3CreN mice (GluN2C^iCre/+^, Grin2CiCre) (Miyazaki et al., 2012). These mice were genotyped using PCR to confirm the presence of the LKB1 flox and Cre recombinase genes.

### Tissue preparation and immunohistochemistry of cerebellum

Mice were deeply anesthetized using either pentobarbital or a mixture of medetomidine hydrochloride (0.75 mg/kg; Zenoaq, Japan), midazolam (4.0 mg/kg; Sandoz, Japan) and butorphanol tartrate (5.0 mg/kg; Meiji Seika Pharma, Japan). Transcranial perfusion was performed with a fixative containing 4% paraformaldehyde (PFA), 10% picric acid in 0.1 M phosphate buffer (PB, pH 7.4). The cerebellum was extracted and cut into 100-µm-thick sagittal sections on a Microslicer (DTK-1000N; DOSAKA EM, Japan). For immunohistochemistry, the cerebellum sections were incubated in a blocking buffer (2% horse serum, 2.5 mg/ml carrageenan, 0.1% NaN_3_, and 0.2% Triton X-100 in PBS), and then incubated in primary antibodies overnight at room temperature. The primary antibodies used were: anti-VGluT1 (1:1000, guinea pig) (Hagiwara et al. 2018), anti-VGluT2 (1:200, guinea pig, Af810; Frontier Institute Co., Ltd., Japan), anti-calbindin (1:1000, mouse, C9848; Sigma-Aldrich, USA), and anti-Iba1 (1:500, rabbit, 019-19741; Fujifilm-Wako, Japan). After washing, sections were incubated with Alexa Fluor-conjugated secondary antibodies (Thermo Fisher Scientific, USA) for 1–2 h at room temperature. Immunolabeled samples were viewed under a confocal laser microscope (LSM900/Airyscan; Carl Zeiss AG, Germany) and analysed using Fiji.

### Tissue preparation and immunohistochemistry of muscles and spinal cord

Hindlimb muscles were collected and weighed. The soleus and gastrocnemius were weighed together. The TA muscle was isolated, and its tendons were pinned to maintain a stretched position. Muscles were fixed in 4% PFA in 0.1 M PB (pH 7.4) for 30 minutes and then cryoprotected by immersion in 20% (w/v) sucrose in PBS at 4 °C. Tissues were embedded in cryoprotectant medium (Neg-50; Thermo Fisher Scientific) for freezing. For cross-sections, the TA muscle oriented vertically on a base tragacanth gum (Fujifilm) was rapidly frozen in isopentane cooled with liquid nitrogen for 30 seconds to 1 minute and then stored at −80 °C. Longitudinal sections (30 µm thick) and cross sections (10 µm thick) were prepared using a cryostat (CM3050S; Leica, Germany) and mounted on glass slides.

Spinal cords were fixed with 4% PFA perfusion, immersed in 30% (w/v) sucrose in PBS at 4 °C, and trimmed. L1–L5 segments were embedded in a cryoprotectant medium. The L4–L5 region was sectioned at a 20 µm thickness and collected on glass slides. The prepared frozen sections were stored at −20°C. The collected sections were post-fixed in 4% PFA in 0.1 M PB (pH 7.4) for 3–5 minutes, washed with 0.1 M PB, and blocked for 30 minutes to 1 hour in blocking solution containing 2.0% (v/v) horse serum, 1.0% Block Ace (Dai Nippon Pharmaceutical, Japan), and 0.2% Triton X-100 in PBS. Sections were then incubated overnight with the following primary antibodies: anti-laminin α2 antibody (1:500, rat, CL3450; Santa Cruz Biotechnology, USA), anti-ChAT antibody (1:100, goat, 4116856; Millipore, USA), and anti-Cre antibody (1:500, mouse, MAB3120; Millipore). After washing, sections were incubated with appropriate Alexa Fluor-conjugated secondary antibodies for 1–2 hours. Images were acquired using a confocal microscope (LSM900/Airyscan) and analyzed with Fiji.

### FT-GO immunofluorescence in neuromuscular junction

Staining of neuromuscular junctions was performed using the multiplex fluorescent tyramide signal amplification system, known as the FT-GO method, as previously described (Yamauchi et al. 2022). Longitudinal sections of the TA were incubated in 1.0% H_2_O_2_ in PBS for 30 minutes to quench endogenous peroxidase activity, followed by washing in 0.3% PBSx (0.3% [v/v] Triton X-100 in PBS). The primary antibody, anti-Synaptophysin antibody (1:200, rabbit, MA5-16402; Thermo Fisher Scientific) diluted in PBS-XCD (0.3% PBSx, 1% [v/v] horse serum, 0.12% [w/v] carrageenan, and 0.02% [w/v] Thimerosal [Nacalai Tesque, Japan]) was incubated overnight. After washing with 0.3% PBSx, the sections were incubated with the secondary antibody, POD F(ab’)_2_ Dk Anti-Rb IgG (1:200, 711-036-152; Jackson Immuno Research, USA) diluted in PBS-XCD for 4 hours. After washing with 0.3% PBSx followed by 0.1 M PB, the FT-GO reaction solution (2.0% BSA in 0.1 M PB, 10 µM CF568 tyramide [Biotium, USA], 3 µg/ml glucose oxidase [Nacalai Tesque], 20 µg/ml β-D-glucose [Nacalai Tesque]) was applied for 30 minutes. After washing with 0.3% PBSx, sections were incubated with Alexa Fluor 488-conjugated α-bungarotoxin (1:500; Thermo Fisher Scientific) diluted in PBS-XCD for 1–2 hours. Finally, the sections were mounted using ProLong Glass Antifade Mounting Medium (Thermo Fisher Scientific) and imaged using a confocal microscope.

### Image Analysis

Cerebellar sections stained with anti-VGlut1 antibody and DAPI were imaged using a tiling method on a confocal microscope. The ML, GL, and WM areas were delineated using the freehand selection tool in Fiji. The Line tool measured ML thickness in lobule IV/V and IX at three distinct points per image. Quantification of cells in the ML and GL was performed by manually counting DAPI-positive cells. For analysis of PC dendrites and CF inputs, Z-stack images of sections immunostained with anti-calbindin and anti-VGlut2 antibodies were acquired. The height of the primary dendrite from the PC body to the pial surface of ML, as well as the height of VGlut2 positive puncta colocalized with the PC dendrite, were measured using the line tool. These values were used to calculate the relative position of CF inputs along the PC dendrites. Microglia quantification was performed using Z-stack images of cerebellar sections stained with anti-Iba1 antibody. The number of Iba1-positive microglia within the ML and GL was manually counted.

For NMJ analysis, Z-stack images were acquired from TA longitudinal sections double-stained with anti-synaptophysin and α-bungarotoxin. The length and width of individual NMJs were measured on the α-bungarotoxin-labeled images using the line tool. The postsynaptic (α-bungarotoxin) and presynaptic terminal (anti-synaptophysin) areas were quantified by binarizing images and applying the analyze particles function, excluding objects smaller than 15 pixels. The innervation ratio was calculated as the ratio of the presynaptic area to the postsynaptic area. Muscle fiber size and density were evaluated using tiling images of anti-laminin labeling. The number and area of individual muscle fibers within 3-5 regions of interest (400 pixels × 400 pixels, 311.96 µm × 311.96 µm) were determined using the analyze particles function after image binarization.

### Western Blotting

The cerebellum and spinal cord were collected from mice anesthetized with isoflurane, and samples were immediately frozen in liquid nitrogen. The tissues were homogenized, and 10 µg of protein from each sample was separated using SDS-PAGE. The separated proteins were then transferred to a PVDF membrane (Millipore). The following primary antibodies were used: anti-LKB1 antibody (1:1000, rabbit, 07-694; Millipore), anti-AMPKα antibody (1:1000, rabbit, 2532; Cell Signaling Technology, USA), anti-Cre antibody (1:1000, rabbit, 69050; Millipore), HRP-conjugated anti-GAPDH antibody (1:1000, rabbit, 3683; Cell Signaling Technology), and anti-Actin antibody (1:5000, mouse, MAB1501; Millipore). For all targets except GAPDH, HRP-labeled secondary antibodies were applied. Chemiluminescent images were captured using an Image Quant LAS4000 system (Cytiva) and analyzed with Fiji.

### Rotarod Test

Motor coordination and balance were assessed using an accelerating rotarod apparatus (Rota-Rod 47600; UGO Basile, Italy) with a rod diameter of 3 cm. Mice were placed individually on the rotating rod, which gradually accelerated from 4 to 40 rotations per minute over a 5-minute period. The latency to fall was recorded for each trial, with a maximum cutoff time of 300 seconds. Each mouse underwent two trials per session, separated by a brief rest period, and the average latency was used as the representative value for that session. Behavioral testing was conducted at the following ages: 4–5 weeks, 8–9 weeks, 12–13 weeks, 16–17 weeks, and 20–21 weeks.

### Markerless tracking of gait and body balance

To assess gait and body balance, mice were allowed to move freely on a transparent acrylic platform (20 cm × 42 cm) while being recorded from below using an RGB-D camera (Azure Kinect DK; Microsoft, USA) for 10 minutes at 30 frames/second. Markerless tracking of body parts was performed using DeepLabCut (Mathis et al. 2018). A total of 500 frames were extracted from the recorded video and manually labeled to identify 10 landmarks: the nose, all four limbs, and five evenly spaced points along the tail. Using the trained DeepLabCut model, mouse movements were tracked in 2D and 3D spaces as previously described (Oikawa et al. 2023; Tsuruda et al. 2023). The following behavioral parameters were quantified from the tracking data: total distance traveled (activity, m), walking speed (mm/second), central zone occupancy (%), inter-limb distance (forelimbs to hindlimbs, mm), distance from hindlimbs to tail0 (mm), projected area formed by the four limbs (mm^2^), nose height (mm), tail midpoint height (tail2, mm), difference in height between the tail0–tail2 (mm). Additionally, behavior states were classified into four categories: Walking, Not Walking, Before Walking, and Motionless. For each mouse, the average or total value of each parameter was calculated per session and compared between control and LKB1 cKO mice at 4–5 weeks, 8–9 weeks, 12–13 weeks, and 16–17 weeks of age.

### Electron microscopy

Mice were perfused with fixative solution containing 2% PFA and 2% glutaraldehyde in 0.1 M PB (pH 7.4), followed by immersion fixation of the brain overnight at 4 ℃. Sagittal sections of the cerebellum (150 μm thick) were washed with 0.1 M PB and then post-fixed in 2% osmium tetroxide in 0.1M PB for 40 minutes. The osmium-fixed sections were then washed with distilled water (dH_2_O) and dehydrated through a graded ethanol series (50%, 70%, 80%, 90%, 95%, and three changes of 100%), with each step lasting 30 minutes. Sections were further dehydrated with a 1:1 mixture of 100% ethanol and propylene oxide for 1 hour and pure propylene oxide two times for 1 hour each.

For resin embedding, Durcupan ACM resin (Fluka, Switzerland) was gradually infiltrated by mixing with propylene oxide in 3:1, 1:1, and 1:3 ratios, each for 12 hours. The tissue was then infiltrated twice with pure resin for 12 hours each, followed by final replacement with fresh resin for 5 hours. Polymerization was carried out at 60 ℃.

Embedded cerebellar sections were trimmed and cut into semi-ultrathin sections (100 nm thick) using an ultramicrotome (Leica EM UC7), and mounted onto glass slides or silicon wafers. Sections were stained with 0.4% (w/v) uranyl acetate followed by lead citrate and coated with osmium using an osmium coater (HPC-1SW) for 1 second, as previously described (Sato et al. 2025). Images were acquired using a field-emission SEM (SU8220; Hitachi High-Tech, Japan) at an accelerating voltage of 5.0 kV with a YAG backscattered electron detector.

Low-magnification images (10k) were acquired from the middle region of the ML to calculate synaptic density. High-magnification images (40k), each containing 1–3 synapses, were obtained for synaptic ultrastructural analysis. The areas of presynaptic terminals and MT were measured using the freehand selection tool, and the length of the PSD was measured using the line tool in Fiji.

### SBF-SEM and 3D reconstruction of PF

Brains were fixed using perfusion of 2% PFA and 2.5% glutaraldehyde in 0.1 M PB followed by immersion overnight in the same fixative at 4 ℃. Sagittal sections of the cerebellum were trimmed to isolate lobule IX (sample dimensions approximately 2–3 mm). The sample was stained and embedded in the resin as previously described with some modifications (Ohno et al. 2014). The tissues were washed four times with PBS for 3 minutes and sequentially incubated with 2% OsO_4_ in 1.5% potassium ferrocyanide for 60 minutes at 4℃, filtered 1% thiocarbohydrazide solution for 20 minutes, 2% OsO4 for 30 minutes at room temperature, 2% uranyl acetate overnight at 4℃, and a 0.66% lead aspartate solution at 50 °C for 2 hours. Samples were washed with double distilled water after each incubation. Then, the tissues were dehydrated in a graded ethanol series (60%, 80%, 90%, and 95% for 5 minutes each), infiltrated with acetone dehydrated with a molecular sieve, a 1:1 mixture of resin (Durcupan; Sigma-Aldrich) and acetone, and 100% resin. Upon embedding, 5% Ketjen black was added to increase the resin conductivity. After curing, the resin blocks were transferred onto rivets for imaging, and approximately 1-μm-thick sections from each block were transferred to glass slides for light microscopy observation with an optical microscope. The block containing the identified lobule was trimmed, and its surface was treated with gold sputtering to increase block conductivity. The block was then imaged with a field-emission SEM (Merlin; Carl Zeiss AG; with Gemini II column and field-emission electron gun) equipped with a 3View Digital Micrograph (version 3.21.1374.0), and OnPoint detector (Gatan, Inc., USA) but not with focal charge compensation. The acceleration voltage was 1.2 kV, and the dwelling time was 0.8 µs/pixel. For 3D reconstruction, 500–600 serial SEM images at a 12-nm/pixel resolution and 50-nm/slice thickness were obtained at two distinct regions of interest at the middle of the ML.

## QUANTIFICATION AND STATISTICAL ANALYSIS

Data are presented as mean ± SEM. Groups were compared using the two-tailed Student’s t-test or two-way analysis of variance with Tukey’s post hoc test. P < 0.05 was considered statistically significant.

## Notes

### Competing Interest Statement

The authors have declared no competing interest.

