## Supplemental Figs for "Depletion of LKB1 in cerebellar granule cells affects parallel fiber–Purkinje cell synaptic organization and causes progressive locomotion deficits"

### Supple Fig. 1

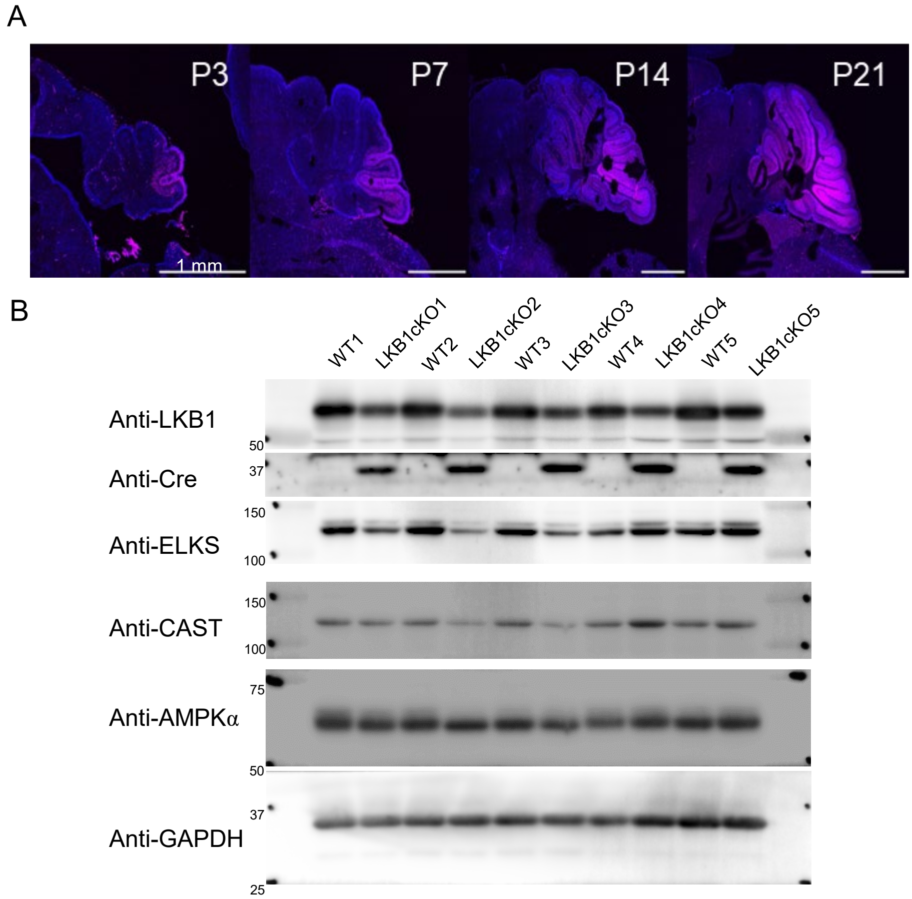


**Supplementary figure 1 Expression of Cre recombinase and loss of LKB1 in the cerebellum of LKB1 cKO mice.** (A) Representative images of Grin2C-iCre;Ai14 mice showing the developmental pattern of Cre recombinase activity. Cre expression was first detected in the posterior cerebellum at postnatal day 3 (P3), expanded progressively from P7 to P14, and by P21 tdTomato expression was observed throughout cerebellar granule cells, indicating widespread Cre recombinase activity. (B) Original western blot showing reduced LKB1 expression accompanied by Cre expression in the cerebellum.

Supple Fig. 2

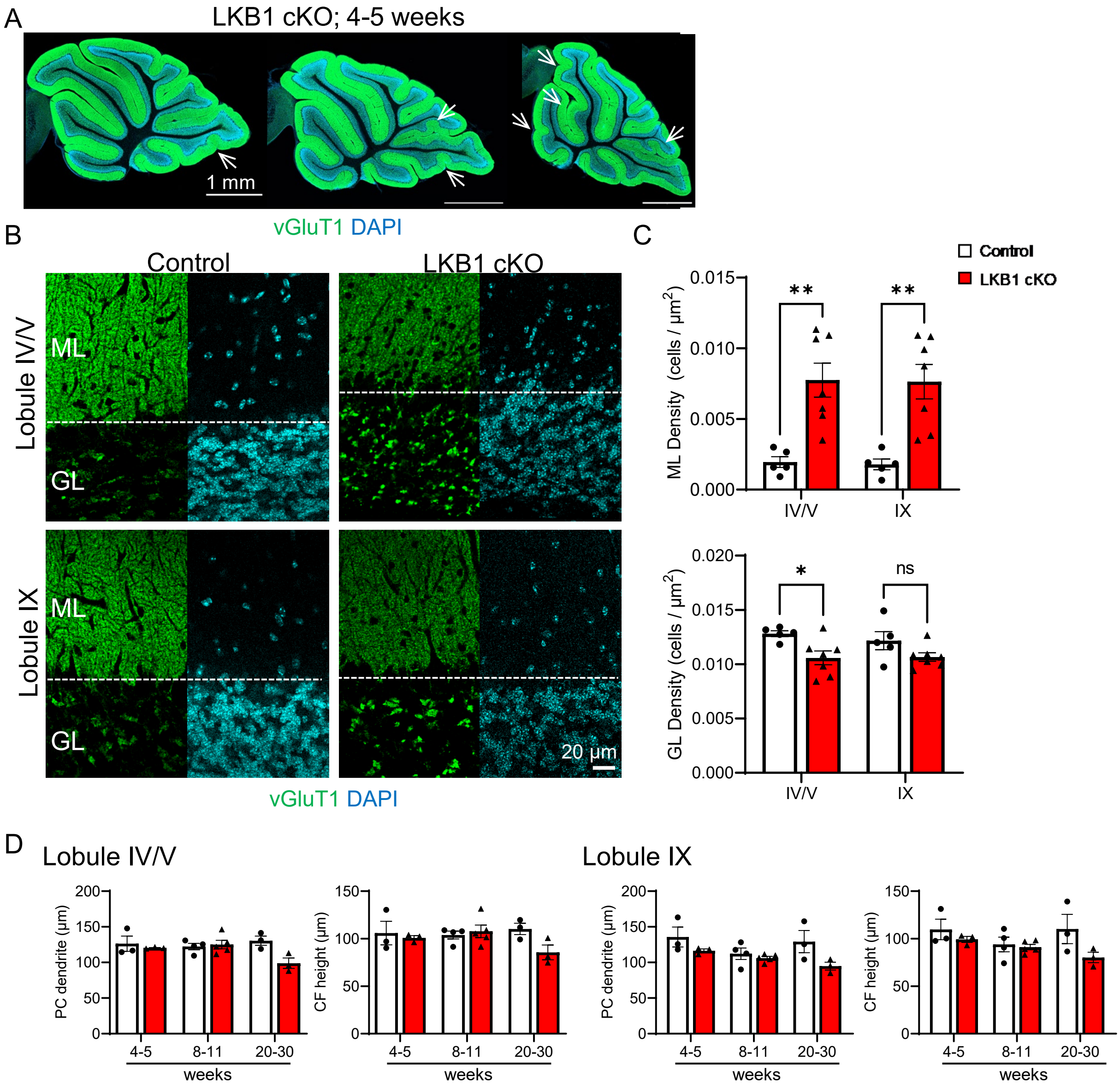


**Supplementary figure 2 Loss of LKB1 in granule cell affects cerebellar morphogenesis**

(A) In 8–10 week LKB1 cKO mice, cerebellar atrophy was predominantly observed in the posterior lobule IX. In contrast, some 4–5 week LKB1 cKO mice exhibited atrophy in the anterior lobules II–IV. The timing of Cre-mediated recombination during development is likely to vary among individual mice. In animals in which Cre expression begins earlier and extends to the anterior cerebellum during early development, cerebellar atrophy may also occur in the anterior region. Conversely, the rarity of anterior cerebellar atrophy in 8–10-week-old mice suggests that animals with earlier Cre expression may die before reaching 10 weeks of age. (B-C) In the molecular layer, DAPI-positive cell density was significantly higher in LKB1 cKO mice than in controls, suggesting that granule cell migration is partially impaired, consistent with previous findings in Atoh1-Cre-mediated LKB1 knockout mice. In contrast, granule cell density showed a trend toward reduction in LKB1 cKO mice. Mean ± SEM, * $P < 0.05$ (Two-way ANOVA with Tukey test). (D) Quantitative analysis of Purkinje cells labeled with anti-calbindin antibody (green) and climbing fibers labeled with anti-VGluT2 antibody (magenta) in Fig. 2D showed that Purkinje cell dendrite length and climbing fiber height were slightly reduced in aged LKB1 cKO mice, although these differences were not statistically significant.

### Supple Fig. 3

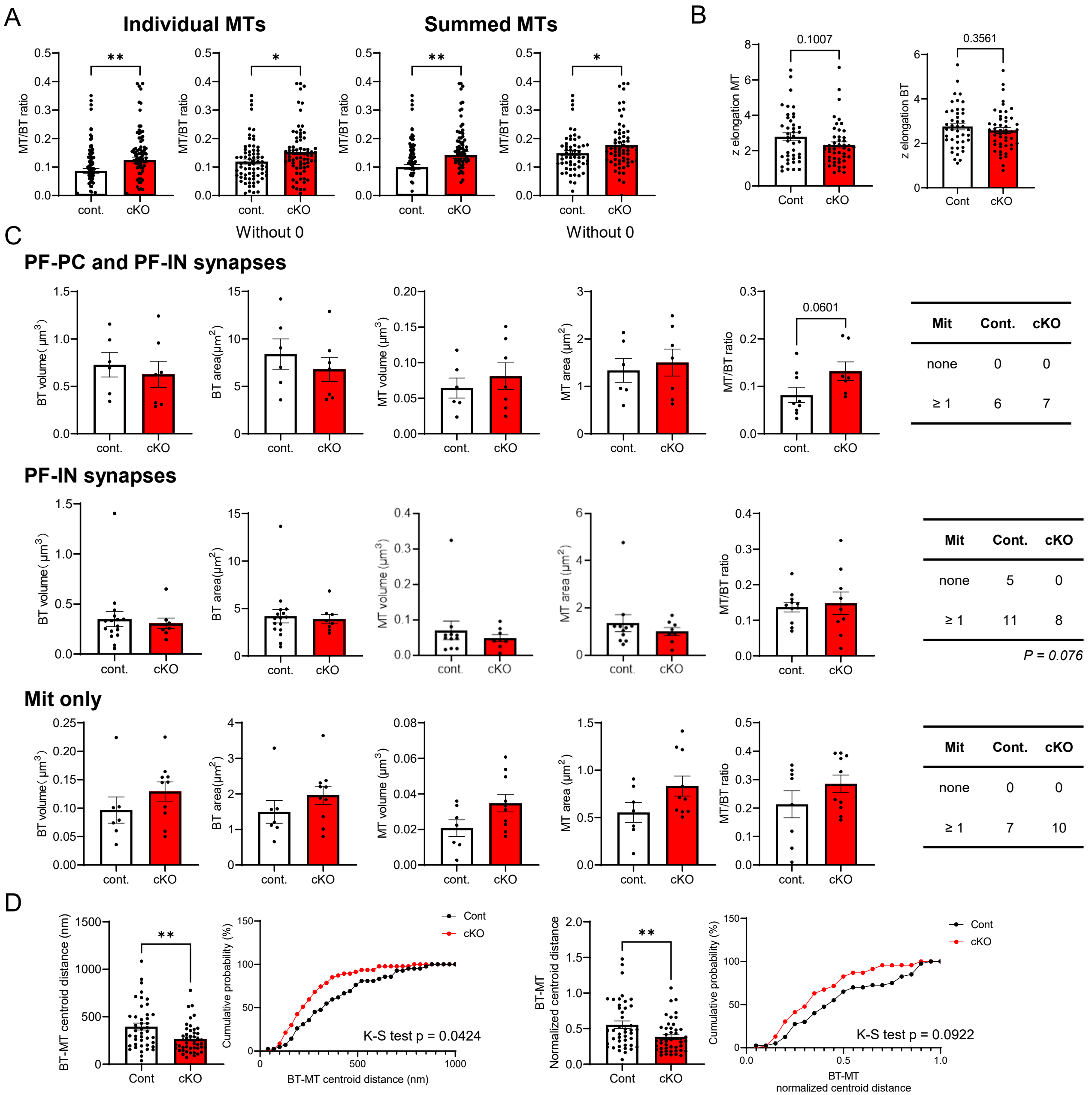


**Supplementary figure 3.** Evaluation of PF BTs and mitochondrial distribution in control and LKB1 cKO mice.

(A) Quantification of the MT/BT volume ratio using individual MT volumes or the summed MT volume per BT. Each analysis was performed either including BTs without MTs (zero values) or excluding BTs without MTs. (B) The Z-elongation ratio of MTs and BTs at PF–PC synapses was calculated as Lz/Dxy, where Lz is the Z-axis length (number of sections × section thickness) and Dxy is the equivalent circular diameter estimated from the reconstructed volume (V) as $2\sqrt{(V/\pi Lz)}$. Higher ratios indicate greater elongation along the Z-axis. (C) Quantitative analysis of PF BTs and MT morphology, including BT volume and surface area, MT volume and surface area, the MT/BT volume ratio, and the distance between adjacent BTs. BTs were classified into three groups: BTs forming synapses with both PC spines and inhibitory neuron (IN) dendrites, BTs forming synapses only with IN dendrites, and BTs without synaptic contacts but containing MT. BTs with PC and IN synapses: Control, n = 6; cKO, n = 7. BTs with IN synapses only: Control, n = 16; cKO, n = 8. BTs without synapses (Mit only): Control, n = 7; cKO, n = 10. Mean ± SEM. (D) Quantitative analysis of MT localization within individual BTs based on the distance between the centroids of BTs and MTs. To account for differences in BT and MT size, centroid distances were normalized to their respective volumes. The cumulative probability distribution was shifted to the left in LKB1 cKO mice, indicating that MT were located closer to the BT center.

### Supple Fig. 4

A

| | Cont | cKO |
|---|---|---|
| spine | 45 | 50 |
| dendrite | 16 | 8 |

Chi-square test p = 0.091

B

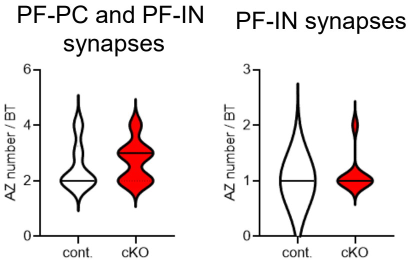


**Supplementary figure 4.** Synaptic connectivity and active zone organization of PF BTs in LKB1 cKO mice.

(A) Quantification of the proportion of BTs forming synapses with PC spines or IN dendrites. LKB1 cKO mice showed a tendency toward a higher proportion of BTs forming synapses with PC spines, although the difference was not statistically significant. (B) Quantification of the number of active zones per BT associated with PC spines or IN dendrites. No significant differences were observed between control and LKB1 cKO mice.

Supple Fig. 5

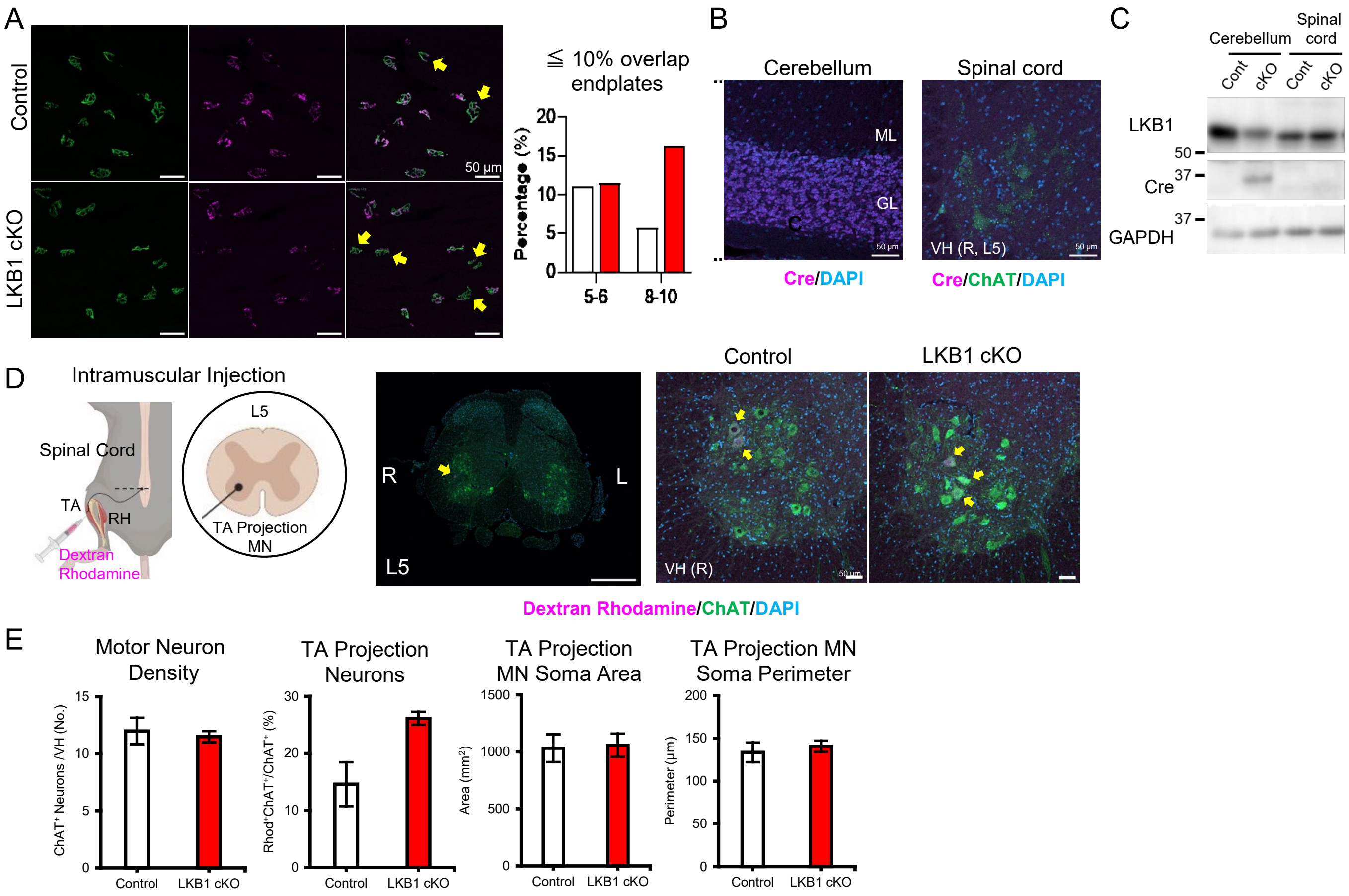


**Supplementary figure 5. Presynaptic innervation of TA-projecting motor neurons**

(A) Representative images showing the overlap between pre- and postsynaptic terminals. Yellow arrows indicate neuromuscular junctions with <10% presynaptic innervation. Quantification was performed on 36–69 neuromuscular junctions per group (Control: 5–6w = 3 mice; 8–10w = 6 mice; LKB1 cKO: 5–6w = 5 mice; 8–10w = 5 mice). (B) In Grin2C-iCre$^{+/-}$ mice, anti-Cre immunostaining revealed robust Cre recombinase expression in cerebellar granule cells, whereas no Cre expression was detected in ChAT-positive motor neurons of the spinal cord. ML, molecular layer; GL, granule cell layer. (C) Western blot analysis confirming LKB1 and Cre expression in the cerebellum (Cb) and spinal cord. (D) Schematic illustration of the retrograde labeling for TA-projecting motor neurons and a representative immunofluorescence image of the L5 spinal cord. Mice were deeply anesthetized with a triple-anesthetic mixture, and approximately 10 μL of tetramethylrhodamine-conjugated dextran (5 mg/mL, Thermo Fisher Scientific) was injected into the TA muscle of the right hindlimb. Spinal cords were collected four days after injection. In the ventral horn (VH), retrogradely labeled TA-projecting motor neurons were visualized by rhodamine fluorescence (magenta), and ChAT-positive motor neurons were identified by immunostaining (green). (E) Quantification of TA-projecting motor neurons were not significantly different between control and LKB1 cKO mice. Control, N = 3; LKB1 cKO, N = 3; Mean ± SEM.

#### Supplementary table 1. Gait analysis

| | Parameters | 4-5w | | | | 8-9w | | | | 12-13w | | | |
|---|---|---|---|---|---|---|---|---|---|---|---|---|---|
| | | Control | | LKB1 cKO | | Control | | LKB1 cKO | | Control | | LKB1 cKO | |
| | | Male | Female | Male | Female | Male | Female | Male | Female | Male | Female | Male | Female |
| Total | Activity (m) | 11.30 | 29.34 | 21.03 | 29.95 | 29.48 | 28.84 | 24.81 | 21.73 | 23.97 | 19.10 | 24.29 | 2.13 |
| | Central Position Rate (%) | 8.79 | 8.69 | 8.42 | 7.07 | 5.47 | 8.38 | 9.09 | 13.24 | 6.79 | 12.07 | 10.50 | 5.35 |
| | Head Direction (mm) | 1.72 | 4.47 | 2.67 | 4.68 | 3.66 | 5.12 | 3.34 | 3.86 | 4.09 | 3.38 | 3.90 | 1.01 |
| | Walking Head Direction (mm) | 3.79 | 6.82 | 4.56 | 6.73 | 5.50 | 6.36 | 5.06 | 6.83 | 6.36 | 7.58 | 6.31 | 1.79 |
| | Walking Speed (mm/s) | 68.11 | 106.92 | 90.20 | 112.71 | 112.29 | 110.10 | 95.97 | 105.97 | 108.39 | 112.14 | 105.18 | 78.89 |
| | Fore-Hind Distance (mm) | 40.67 | 37.76 | 36.34 | 37.57 | 43.72 | 34.26 | 41.61 | 32.71 | 39.99 | 35.42 | 32.33 | 30.10 |
| | Hind-Tail0 Distance (mm) | 16.02 | 14.82 | 21.47 | 10.25 | 18.50 | 15.78 | 20.76 | 21.14 | 18.98 | 13.25 | 23.98 | 26.70 |
| | Limbs Area ($mm^2$) | 1025.76 | 709.51 | 655.10 | 676.08 | 766.71 | 610.77 | 838.22 | 624.95 | 795.35 | 727.05 | 678.86 | 579.65 |
| | Height of Tail (mm) | 36.19 | 36.65 | 39.97 | 34.78 | 39.81 | 37.13 | 48.48 | 41.62 | 39.01 | 38.99 | 46.38 | 38.55 |
| | Height of Tail0-Tail2 (mm) | 22.02 | 20.87 | 24.54 | 20.70 | 23.33 | 21.72 | 31.26 | 26.02 | 22.14 | 23.30 | 31.37 | 23.12 |
| | Height of Nose (mm) | 12.22 | 15.28 | 12.66 | 15.81 | 14.79 | 17.08 | 14.99 | 13.55 | 16.16 | 16.82 | 11.31 | 14.45 |
| Walking | Fore-Hind Distance (mm) | 37.81 | 42.21 | 35.78 | 49.91 | 46.06 | 42.89 | 46.37 | 40.85 | 45.11 | 39.34 | 42.44 | 47.47 |
| | Hind-Tail0 Distance (mm) | 28.90 | 12.93 | 17.32 | 9.01 | 17.19 | 13.81 | 20.98 | 19.59 | 16.52 | 13.52 | 23.65 | 22.42 |
| | Limbs Area ($mm^2$) | 875.20 | 871.10 | 733.26 | 923.76 | 824.22 | 798.62 | 926.10 | 782.94 | 919.35 | 806.78 | 908.14 | 1066.88 |
| | Height of Tail (mm) | 36.65 | 42.97 | 43.17 | 40.73 | 43.78 | 42.84 | 53.85 | 50.17 | 45.73 | 45.32 | 54.79 | 56.92 |
| | Height of Tail0-Tail2 (mm) | 22.87 | 24.83 | 25.93 | 24.00 | 26.10 | 25.43 | 34.92 | 30.87 | 27.39 | 27.80 | 37.30 | 39.31 |
| | Height of Nose (mm) | 10.68 | 13.35 | 11.73 | 14.94 | 14.10 | 15.85 | 14.56 | 12.44 | 15.57 | 14.99 | 10.79 | 10.51 |
| Not Walking | Fore-Hind Distance (mm) | 39.11 | 35.66 | 35.23 | 32.13 | 43.19 | 30.88 | 39.67 | 30.33 | 38.60 | 33.72 | 28.60 | 29.79 |
| | Hind-Tail0 Distance (mm) | 15.37 | 15.63 | 21.40 | 10.03 | 18.96 | 16.64 | 21.19 | 21.35 | 19.80 | 13.63 | 24.05 | 26.78 |
| | Limbs Area ($mm^2$) | 978.89 | 620.20 | 611.20 | 572.24 | 745.41 | 537.35 | 799.33 | 584.02 | 763.53 | 684.63 | 594.77 | 570.86 |
| | Height of Tail (mm) | 35.78 | 33.80 | 38.79 | 32.26 | 37.65 | 34.90 | 46.09 | 38.86 | 37.01 | 37.68 | 43.44 | 38.22 |
| | Height of Tail0-Tail2 (mm) | 21.58 | 19.14 | 23.87 | 19.11 | 21.81 | 20.28 | 29.72 | 24.36 | 20.60 | 22.56 | 29.30 | 22.83 |
| | Height of Nose (mm) | 12.42 | 16.13 | 12.95 | 16.22 | 15.13 | 17.54 | 15.34 | 13.85 | 16.36 | 17.09 | 11.48 | 14.52 |
| Motionless | Fore-Hind Distance (mm) | 39.14 | 36.35 | 35.05 | 31.74 | 43.69 | 30.90 | 40.26 | 30.04 | 39.46 | 33.45 | 27.89 | 29.71 |
| | Hind-Tail0 Distance (mm) | 15.54 | 15.50 | 21.60 | 9.36 | 18.46 | 15.98 | 21.15 | 21.24 | 19.55 | 13.33 | 23.88 | 27.81 |
| | Limbs Area ($mm^2$) | 977.89 | 621.96 | 599.57 | 562.87 | 738.19 | 526.05 | 805.99 | 575.28 | 788.52 | 676.40 | 571.33 | 562.99 |
| | Height of Tail (mm) | 35.70 | 32.90 | 38.63 | 31.10 | 37.14 | 33.71 | 45.18 | 38.12 | 36.47 | 37.26 | 43.17 | 38.36 |
| | Height of Tail0-Tail2 (mm) | 21.46 | 18.54 | 23.85 | 18.34 | 21.51 | 19.40 | 29.16 | 23.92 | 20.26 | 22.44 | 29.27 | 22.78 |
| | Height of Nose (mm) | 12.39 | 16.19 | 12.98 | 16.17 | 14.98 | 17.51 | 15.44 | 13.73 | 16.25 | 17.21 | 11.61 | 14.60 |
| Before Walking | Fore-Hind Distance (mm) | 52.60 | 37.35 | 33.30 | 37.29 | 44.87 | 35.35 | 42.27 | 33.62 | 39.82 | 33.44 | 35.94 | 44.59 |
| | Hind-Tail0 Distance (mm) | 30.17 | 14.86 | 17.52 | 11.48 | 20.29 | 17.27 | 20.07 | 19.19 | 18.66 | 18.35 | 24.69 | 12.70 |
| | Limbs Area ($mm^2$) | 1360.54 | 693.61 | 713.39 | 634.86 | 791.83 | 617.12 | 850.68 | 616.71 | 720.81 | 668.31 | 788.92 | 1230.05 |
| | Height of Tail (mm) | 37.76 | 36.90 | 37.95 | 35.38 | 37.68 | 36.93 | 46.24 | 41.18 | 39.23 | 39.51 | 42.76 | 41.85 |
| | Height of Tail0-Tail2 (mm) | 25.40 | 22.18 | 23.97 | 21.52 | 22.40 | 22.01 | 30.08 | 26.43 | 23.18 | 24.91 | 29.21 | 27.44 |
| | Height of Nose (mm) | 13.18 | 15.78 | 13.01 | 17.38 | 16.52 | 17.89 | 15.90 | 14.87 | 17.74 | 17.60 | 12.04 | 12.12 |

#### Supplementary table 2. Grip Strength

| Male | Contol | | | LKB1 cKO | | |
|---|---|---|---|---|---|---|
| | 4-5w | 8-9w | 12-13w | 4-5w | 8-9w | 12-13w |
| Grip Strength (g) | 131.42 | 173.50 | 181.89 | 122.92 | 147.83 | 180.50 |

| Female | Contol | | | LKB1 cKO | | |
|---|---|---|---|---|---|---|
| | 4-5w | 8-9w | 12-13w | 4-5w | 8-9w | 12-13w |
| Grip Strength (g) | 180.72 | 176.20 | 143.47 | 181.58 | 184.83 | 123.00 |
